# DNA methylation affects gene expression but not global chromatin structure in *Escherichia coli*

**DOI:** 10.1101/2025.01.06.631547

**Authors:** Willow Jay Morgan, Haley M. Amemiya, Lydia Freddolino

**Affiliations:** Department of Biological Chemistry, University of Michigan Medical School, Ann Arbor, MI 48109, USA; Cellular and Molecular Biology Program, University of Michigan Medical School, Ann Arbor, MI 48109, USA; MOMA Therapeutics, Cambridge MA 02140; Department of Computational Medicine and Bioinformatics, University of Michigan Medical School, Ann Arbor, MI 48109, USA

## Abstract

The activity of DNA adenine methyltransferase (Dam) and DNA cytosine methyltransferase (Dcm) together account for nearly all methylated nucleotides in the *Escherichia coli* K-12 MG1655 genome. Previous studies have shown that perturbation of DNA methylation alters *E. coli* global gene expression, but it is unclear whether the methylation state of Dam or Dcm target sites regulates local transcription. In recent genome-wide experiments, we observed an underrepresentation of Dam sites in transcriptionally silent extended protein occupancy domains (EPODs), prompting us to hypothesize that EPOD formation is caused partially by low Dam site density. We thus hypothesized that a methylation-deficient version of MG1655 would show large-scale aberrations in chromatin structure. To test our hypothesis, we cloned methyltransferase deletion strains and performed global protein occupancy profiling using high resolution *in vivo* protein occupancy display (IPOD-HR), chromatin immunoprecipitation for RNA polymerase (RNAP-ChIP), and transcriptome abundance profiling using RNA-Seq. Our results indicate that loss of DNA methylation does not result in large-scale changes in genomic protein occupancy such as the formation of EPODs, indicating that the previously observed depletion of Dam sites in EPODs is correlative, rather than causal, in nature. However, loci with dense clustering of Dam methylation sites show methylation-dependent changes in local RNA polymerase and total protein occupancy, but local transcription is unaffected. Our transcriptome profiling data indicates that deletion of *dam* and/or *dcm* results in significant expression changes within some functional gene categories including SOS response, flagellar synthesis, and translation, but these expression changes appear to result from indirect regulatory consequences of methyltransferase deletion. In agreement with the downregulation of genes involved in flagellar synthesis, *dam* deletion is characterized by a swimming motility-deficient phenotype. We conclude that DNA methylation does not control the overall protein occupancy landscape of the *E. coli* genome, and that observable changes in gene regulation are generally not resulting from regulatory consequences of local methylation state.

**IMPORTANCE:** Previous studies of *E.* coli chromatin structure revealed a statistical association between the presence of silenced, highly protein occupied regions of the genome and depletion of modification sites for Dam methyltransferase. Here, we show that loss of DNA methylation does not substantively affect global chromatin structure in *E. coli*, thus demonstrating that the previously observed correlation was not causal. However, we observed specific methylation-dependent changes in gene expression, particularly affecting the SOS response, flagellar synthesis, and translation. These effects appear to be indirect regulatory consequences of methyltransferase deletion. Our work clarifies the role of methylation in chromatin structure and regulation, providing new insights into the mechanistic basis of gene expression and chromatin structure in *E. coli*.

## INTRODUCTION

DNA methylation in bacteria has well-supported roles in phage defense, chromosomal replication, DNA repair, and the regulation of gene expression^1^. Most examples of bacterial methyltransferases are involved in restriction-modification phage defense systems, which involve the methylation of target sequences in bacterial DNA to protect against the endonuclease activity of restriction enzymes which are synthesized by the bacteria to cleave non-methylated phage DNA^2,3^. Methylated DNA generally results from the activity of methyltransferases which cleave the methyl group from S-adenosyl-L-methionine (SAM) and transfer it to adenine or cytosine^4,5^. DNA methyltransferases have been found a wide range of bacterial species to be involved in chromosomal maintenance, replication, DNA repair, cell cycle control, and virulence^6–13^. There are also some examples of bacterial DNA methylation influencing local binding of regulatory proteins in promoters which impacts transcription^14–19^, but it is unclear whether this is a common phenomenon in *Escherichia coli* ^1,20–23^. In the case of commonly studied *E. coli* K-12 laboratory strains, two major methyltransferases are known: Dam and Dcm.

DNA adenine methyltransferase (Dam) in *Escherichia coli* catalyzes the formation of N^6^-methyladenine in the target motif 5’-G<u>A</u>TC-3’^24^. Dam is conserved within Enterobacteriaceae. Over 99% of methylated adenines at ∼19,000 sites in the genome of the *E. coli* K-12 strain MG1655 result from Dam activity^1^. Dam is characterized as an orphan methyltransferase^25^ as there is no known cognate restriction enzyme that cleaves at Dam methylation sites in *E. coli* K-12. Dam methylation has been implicated as a critical element of chromosomal maintenance, as the methylation state of ∼11 Dam target sites at the origin of replication (oriC) regulates initiation of chromosomal replication^26–28^. During replication, Dam sites throughout the chromosome are processively methylated whilst lagging the replication fork, producing a transient hemi-methylated state utilized by DNA repair machinery to differentiate between the template and newly-synthesized strands in the event that mismatch repair is needed^29–32^.

*E. coli* DNA cytosine methyltransferase (Dcm) methylates the inner cytosine in its 5’-C<u>C</u>WGG-3’ target motif^33^, and Dcm is conserved within *Escherichia*. Dcm appears to be responsible for all cytosine methylation at ∼12,000 sites in the *E. coli* K-12 genome^1,22^. Taken together with Dam, these two methyltransferases produce nearly the entire *E. coli* K-12 methylome^1,34^. Dcm is known to have a cognate restriction enzyme, *Eco*RII, which is not found in K-12 strains^35^. The role of Dcm methylation in other cellular functions is less well-characterized than in the case of Dam. While there are few to no changes in growth dynamics when *dcm* is deleted, there does appear to be a fitness benefit associated with Dcm methylation in long-term stationary phase^22,36^. Dcm is also involved in Very Short Patch (VSP) repair in *E. coli* K-12, where the repair-associated endonuclease Vsr nicks double-stranded DNA at the Dcm target motif when the inner methylated cytosine is deaminated to thymine (5’-C**T**WGG-3’)^37^. It is unclear whether the methylation state of the cytosine residue impacts this VSP repair process.

Perturbation of DNA methylation alters *E. coli* global gene expression to some extent, but the mechanisms by which the methylation state of Dam or Dcm sites regulates local transcription are not fully understood^20–22,38,39^. In one example of DNA methylation acting as a transcriptional regulator, two nucleoid-associated proteins (NAPs) and Dam compete for binding to the promoter of the virulence-associated *pap* operon in uropathogenic *E. coli*^16,17^. NAPs are promiscuous DNA-binding proteins that confer chromosomal structure and act as global regulators^40,41^. There are other characterized examples of the methylation state at Dam motifs in NAP binding sites influencing NAP binding affinity – and in some cases, gene expression – across different bacterial strains^11,18,19,42^. A potential mechanism is that DNA methylation-dependent alterations of DNA-protein interactions result from the protrusion of the methyl group into the major groove producing DNA curvature^39,43,44^.

The full extent to which DNA methylation altering NAP occupancy contributes to gene expression changes in *E. coli* K-12 is unknown. A recent study analyzing total protein occupancy data – produced by high resolution *in vivo* protein occupancy display (IPOD-HR) – reported an underrepresentation of Dam target motifs in extended protein occupancy domains (EPODs)^45,46^. EPODs are ≥1 kilobase regions of the genome that have a continuously high protein occupancy signal; EPODs can be considered functional analogs to eukaryotic heterochromatin as EPODs are primarily formed by dense clusters of NAPs that coat DNA and silence local transcription^46,47^.

Our observation that Dam sites are underrepresented in EPODs – in addition to the regulatory cross-talk demonstrated with the *pap* operon and other examples – led us to speculate that there is a causal relationship between DNA methylation state and protein occupancy which contributes to the formation of EPODs in *E. coli* K-12 MG1655. We thus hypothesized that a methylation-deficient version of MG1655 would show large-scale aberrations in chromatin structure (in particular, the formation and locations of EPODs) which might alter the regulation of silenced wild-type genomic regions. To test for such changes, we cloned single deletion mutants of Dam and Dcm (*Δdam* and *Δdcm*, respectively) and a double deletion mutant of both Dam and Dcm (*Δdam/Δdcm*), and we performed global protein occupancy profiling (using the IPOD-HR method^46^) and transcriptome abundance profiling (using RNA-Seq) on these strains to produce global protein occupancy profiles and identify EPOD locations. Our results indicate that, relative to wild-type cells, DNA methylation-deficient mutants of *E. coli* K-12 MG1655 are not characterized by large-scale changes in genomic protein occupancy such as the formation of EPODs. Thus, the reduced abundance of Dam sites in EPODs does not cause EPOD formation (at least, not through reduced methylation density), but rather, is likely the consequence of some shared feature of these regions. However, we have identified a small number of loci with dense clustering of Dam methylation sites for which our data shows methylation-dependent changes in local RNA polymerase and total protein occupancy. Our transcriptome profiling data indicates that deletion of *dam* and/or *dcm* results in significant expression changes within some functional gene categories including SOS response, flagellar synthesis, and translation, but these expression changes appear to result from indirect regulatory consequences of methyltransferase deletion rather than being due to perturbation of interactions between DNA methylation and regulatory proteins at gene promoters. As such, there are no changes in local transcription associated with the dense clusters of Dam sites. Dam deletion mutants were, however, characterized by a swimming motility-deficient phenotype which is likely associated with the downregulation of genes involved in flagellar synthesis. Thus, we find that DNA methylation does not control the overall protein occupancy landscape of the *E. coli* genome, and that changes in gene regulation are generally an indirect effect of loss of Dam methylation, rather than a direct regulatory consequence of local methylation state.

## METHODS

### Bacterial strain construction

The “WT” parental strain of *Escherichia coli* K-12 MG1655 was obtained from Dr. Haley Amemiya, who sourced it from Hani Goodzari (Tavazoie Lab, then at Princeton University) in 2009 as described in Amemiya *et al.*, 2022^47^. This “WT” isolate is isogenic with ATCC 700926 except for an IS1 insertion in *dgcJ* ^47,48^.

*Δdam, Δdcm*, and *Δdam/Δdcm* strains were constructed from the parental “WT” via P1 transduction of a FRT-flanked *kanR* marker from corresponding knockout strains in the Keio collection^49,50^. The *kanR* marker was excised through electroporating the pCP20 helper plasmid – which encodes for Flp recombinase – leaving a small scar in place of the indicated genes’ original open reading frames^51^. Isolated transformants were grown overnight at 42℃ to remove the temperature-sensitive pCP20, and these overnight cultures were non-selectively purified on LB plates grown overnight at 37℃. Candidate colonies were replica plated onto LB and selective plates to confirm the loss of the *kanR* marker and pCP20 plasmid. Sanger sequencing verified the deletion of the indicated gene with the replacement of a small scar.

*ΔlrhA* and *Δdam/ΔlrhA* strains were constructed through P1 transduction of FRT-flanked *kanR* from corresponding Keio collection strain followed by pCP20-mediated recombination as described above.

All constructs were verified using Sanger sequencing through AZENTA Life Sciences GENEWIZ^®^.

### Media and culture conditions

LB (Lennox) media (10 g/L tryptone, 5 g/L yeast extract, 5 g/L NaCl) was used for the above cloning, recovery of cryogenically-preserved *E. coli* cells, and culturing for motility assays. 15 g/L bacteriological agar was added for plates.

MOPS-RDM corresponding to the fully supplemented version of MOPS defined medium in Neidhardt *et al.*, 1974^52^ (with 0.4% glucose as a carbon source) was used to grow *E. coli* cells for IPOD-HR, motility assays, and RNAP-ChIP. 15 g/L bacteriological agar was added to the MOPS-RDM recipe to make plates. Minimal MOPS media was made as specified in Neidhardt *et al.*, 1974^52^ using 0.2% w/v glucose as a carbon source.

PYE (Peptone Yeast Extract; 2 g/L peptone, 1 g/L yeast extract, 1mM MgSO_4_, pH 6.0) was used to grow *Caulobacter crescentus* cells to produce a spike-in reference for IPOD-HR. 20 g/L bacteriological agar was added for plates.

### Cell growth and harvest for IPOD-HR

Our procedures for IPOD-HR largely follow those described in Amemiya *et al*., 2022^47^. Cryogenically preserved cells were streaked onto a plate and isolated colonies were subsequently grown in the same media as used for plating (MOPS-RDM for *E. coli*, PYE for *C. crescentus*) overnight at 37°C with shaking at 200 rpm. The culture was back-diluted into fresh, prewarmed media to an OD600 of 0.003 the next day. The culture was grown to the target OD600 of 0.2 and a 500 µL aliquot (for RNAseq) was taken and added to 1 mL of RNAprotect Bacteria Reagent (Qiagen, Hilden, Germany) and preserved according to the manufacturer’s instructions. The remainder of the culture was treated with a final concentration of 150 μg/ml rifampicin and returned to the same growth conditions for another 10 minutes. The cultures were then rapidly pipetted into 50-mL conical tubes and mixed with concentrated formaldehyde/sodium phosphate (pH 7.4) buffer sufficient to yield a final concentration of 10 mM NaPO4 and 1% w/v formaldehyde.

Crosslinking proceeded for 5 min at room temperature with 300 rpm shaking, and then quenched with an excess of glycine (final concentration 0.333 M) for 10 min with 300 rpm shaking at room temperature. Cells were then chilled on ice for 10 minutes and then pelleted and washed twice with ice-cold phosphate-buffered saline (PBS). The resulting pellets were dried by pipetting residual liquid, and the tubes were then snap-frozen in a dry ice-ethanol bath before being stored at −80°C.

### Cell lysis and DNA preparation

When resuspending frozen cell pellets, two pellets (taken from a single biological replicate) of each sample were separately resuspended with spike-in, sonicated, and then combined into one tube immediately after sonication. Individual frozen *C. crescentus* spike-in cell pellets were resuspended in 600 µL of 1x IPOD lysis buffer (10 mM Tris HCl, pH 8.0; 50 mM NaCl) containing 1x protease inhibitors (Roche Complete Mini, EDTA free, Roche Diagnostics GmbH, Mannheim, Germany) and 1.5 µL of ready-lyse (Epicentre, Madison, WI). The spike-in resuspension was used to resuspend one of the sample cell pellets, and then the resuspended cells were incubated for 15 minutes in a 30°C water bath. Sonication was then performed on all samples using a Branson digital sonifier with a microtip at 25% amplitude for four pulses of 5 seconds with a 5 second rest between each pulse; samples were kept in an ice/water bath during sonication. The two separate tubes for each biological sample were then combined.

DNA digestion was performed by adding to the sonicated lysates 120 μg RNase A (Thermo Fisher Scientific, Waltham, MA), 12 μL DNase I (Fisher product #89835), 10.8 μL 100 mM MnCl2, and 9 μL 100 mM CaCl2, and then incubating on ice for 30 minutes. The digestion was quenched with 100 μL of 500 mM EDTA (pH 8.0), followed by clarification by centrifuge for 10 minutes at 13,000 rpm at 4°C. Aliquots were taken from the clarified lysate for IPOD-HR interface extraction, RNA polymerase chromatin immunoprecipitation, and cross-linking reversal and recovery of DNA as previously described^46^. The procedures described in that reference were replicated here, apart from all the 2-minute centrifuging steps instead being done in 4 minutes.

For DNA recovery, standard phenol-chloroform extraction as ethanol precipitation as described in Ausubel F, 1998^53^, and the dried DNA was resuspended in 100 μL of TEe (10 mM Tris pH 8.0; 0.1 mM EDTA pH 8.0) for input samples, 20 μL of TEe for RNAP-ChIP samples, and 50 μL of TEe for IPOD samples.

### RNA isolation and sequencing preparation

RNA pellets were removed from −80°C and then immediately resuspended in 100 μL TE buffer (10 mM Tris pH 7.5; 1 mM EDTA). The resuspended pellet was treated with 1 μL lysozyme (Ready-Lyse; Lucigen, Ltd.) and incubated for 10 minutes at 4°C, followed by treatment with 10 μL proteinase K and incubation for 10 minutes at room temperature with vortexing every 2 minutes. RNA was then isolated using a Zymo RNA Clean and Concentrate 5 Kit twice for each sample, with a DNase digestion (25 μL eluate from first Zymo clean-up, 58 μL nuclease-free water, 10 μL 10X DNase Reaction Buffer, 2 μL RNase inhibitor, 5 μL Baseline Zero DNase) at 37°C for 30 minutes in-between Zymo kit clean-ups. Samples were then ribo-depleted using a NEBNext rRNA Depletion (Bacteria) Kit according to manufacturer’s instructions – with the exception that 10 μL of RNA sample, containing 1 μg of RNA, was used for probe hybridization. Following ribo-depletion, samples were again cleaned-up with the Zymo Clean and Concentrate 5 Kit and then prepared for sequencing using the NEBNext Ultra II Directional RNA Sequencing Kit and then sequenced as described below for DNA samples.

### Preparation of next-generation sequencing (NGS) libraries

DNA samples were prepared for Illumina sequencing using NEBNext Ultra II Library Prep Kit (NEB product #E7103) and NEBNext Muliplex Oligos for Illumina (96 reactions, NEB product #E6442S). Deviations from manufacturer’s directions to account for low average fragment sizes are described in Freddolino *et al.*, 2021^46^. All libraries were sequenced on an Illumina NextSeq instrument.

Analysis of NGS data, read quality control and preprocessing, DNA sequencing and protein occupancy calling, and feature calling were performed as previously described in Freddolino *et al.*, 2021^46^. However, we used here a more recent version of the IPOD-HR pipeline `ipod_v2.5.7` which can be obtained from https://github.com/freddolino-lab/ipod (matching commit e2c2889 in that repository). Some of the software used in IPOD-HR version 2.5.7 includes: `cutadapt v3.5`^54^, `trimmomatic v0.39`^55^, `bowtie2 v.2.4.4`^56^, and `samtools v1.1.4`^57^; definition files for building a singularity container exactly matching our workflow are available on the github repository noted above.

A summary of the changes introduced between the IPOD-HR version utilized in Freddolino *et al*., 2021^46^ and the verison utilized here are summarized as follows: Following quantile normalization, each replicate of each data type (IPOD, ChIP, input DNA) was median normalized to 100. A pseudocount of 0.25 was then added to each datum. Log_2_ ratios of IPOD or ChIP data relative to input data were calculated for each set of paired replicates. The log_2_ ratios were converted to robust z-scores and log_10_ p-values for visualization as described in Freddolino *et al*., 2021^46^. 95% confidence limits and mean estimates were calculated for log_2_ ratios, log_10_ p-values, and robust z-scores using jackknife sampling of the scores for all three biological replicates of each data type.

EPOD calling was performed similar to as described in Freddolino *et al*., 2021^46^ with the following deviations: EPOD seed regions were identified as any region at least 1,024 basepairs in length over which the median of a 768 bp rolling median exceeded the overall 90^th^ or 75^th^ percentile in the case of strict or loose EPOD calling, respectively, of a 256 bp rolling median over the entire chromosome. EPODs from all biological replicates of each given condition were “merged” into single, contiguous genomic intervals to assess the degree to which EPODs from replicate conditions overlap. EPODs were called separately at the biological replicate level, and then EPOD locations with low reproducibility were dropped from analysis based on an upper limit of 0.05 for the irreproducible discovery rate^58^.

### Methylation motif flagging

To characterize the occupancy changes observed across genotypes relative to the methyltransferase target sites, we scanned each base pair of the *E. coli* U00096.3 genome and assigned each base pair a motif flag. Methyltransferase target sites were identified based on whether a sequence of base pairs matched the target motif of each methyltransferase: “GATC” for Dam, “CCTGG” or “CCAGG” for Dcm.

Scanning for these motifs was done using the `motifs` package from `Biopython`^59^. The output of this `python v3.10.2` script was a listing of each methyltransferase target motif location in the genome. This information was used to create a file containing a list of every single base pair in the genome accompanied by an appropriate motif flag indicating membership to a methyltransferase target site.

### IPOD-HR and RNAP-ChIP occupancy at individual and clustered methylation sites

To capture more of the genomic context surrounding methylation sites, the `slop` command from `bedtools v2.30`^60^ was used to add 50 bp extensions to the start and end positions of each methylation site feature. The IPOD-HR and RNAP-ChIP occupancy scores at the extended methylation site features were found using `bedtools intersect`. Violin plots of the occupancy scores at methylation sites were made using `seaborn v0.11.2`^61^.

The density of methylation sites at genomic loci was determined by counting the number of methylation sites within each extended methylation site, and this count was then added as a flag to the extended methylation site.

Occupancy subtractions between genotypes were done to highlight occupancy changes unique in mutants relative to wild-type. To subtract occupancy between genotypes, negative values were adjusted to “0” for all genotypes and then wild-type occupancy was subtracted from each mutant occupancy track.

### Read end analysis

To identify the read ends of each input sample, `bedtools genomecov -ibam` with the `-5` and `-3` arguments was called on each BAM file output by the IPOD-HR alignment pipeline with the `-bg` argument used to produce bedgraph files. The 5’ and 3’ read ends were then combined into one file for each sample and the read end count at each position was normalized by the total number of million read ends within each sample. 100 basepair flanks were added to each end of the “7 Dam Site” cluster 100 basepair windows and `bedtools intersect` was used to find the normalized read end counts at these dense Dam site clusters. Heatmaps were generated using `seaborn`.

### EPOD analysis

Symmetrized overlap distance was calculated as shown in Equation 1 and as previously described in Amemiya *et al.*, 2022^47^. Overlapping “strict” and “loose” EPODs were found using `bedtools intersect`. The frequency of Dam or Dcm sites in EPODs for each condition was calculated using `bedtools intersect` and then divided by the genomic total frequency of Dam or Dcm sites, respectively.

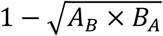

**Equation 1**: Calculation of Symmetrized Overlap Distance, which quantifies the overlap between two EPOD sets *A* and *B*, as previously described in Amemiya *et al.*, 2022^47^. *X_Y_* is the fraction of condition *Y*’s strict EPODs which are genomically overlapped (in at least 1 basepair position) with the loose EPODs in condition *X*.

For analysis of the representation of Dam sites in EPODs while controlling for the AT% of EPODs, each strict EPOD called for wild-type (“genomic EPODs”) was assigned to one of 10 evenly populated bins that were discretized based on AT content (AT%). 1000 random shufflings of the EPOD genomic locations (“shuffled EPODs”), allowing the shuffled locations to overlap original EPOD locations (“overlaps”) or not (“no overlaps”), were produced using `bedtools shuffle`. The shuffled EPODs were then assigned to the AT% bins, and within each AT% bin the Dam sites per kilobase for shuffled EPODs and genomic EPODs were compared by Poisson regression using `R v4.1`^62^ and plotted with `ggplot2 v3.3.5`^63^.

### RNAseq analysis

The NGS data produced from Illumina sequencing of the RNA samples was processed through the IPOD-HR pipeline as described above up to but not including the alignment step. The `Rockhopper v2.0.3` RNAseq analysis system^64–66^ was then utilized to align the processed RNAseq reads to the U00096.3 genome and identify transcripts. Rockhopper returns q-values which represent the statistical significance of differential expression of each transcript between conditions. The log_2_-fold change in transcript values were calculated, with directionality assigned based on whether mutant transcript value was greater (positive) or less (negative) than the wild-type transcript value; plots were generated using `matplotlib v3.5.1`^67^.

### iPAGE

We applied the iPAGE software previously described^68^ to perform gene set enrichment analysis. To produce the data for running iPAGE, the q-values from the Rockhopper analysis described above which compared the transcripts between our mutant and wild-type genotypes were log_10_-transformed and assigned directionality (positive or negative) based on whether the transcript is higher (positive) or lower (negative) in abundance in the mutant genotype relative to wild-type. The directional log_10_q-values were fed into iPAGE as a continuous variable, and so iPAGE created equally populated discrete bins to rank the directional log_10_q-values and calculate their representation within each bin for all transcripts associated with a given GO term.

### Motility regulon expression analysis

The regulons of each regulator of *flhDC* were identified using the transcription factor to gene pairing database reported through RegulonDB^69^. Expression changes between each of the methyltransferase mutants and wild-type were found using the log_2_-ratios of Rockhopper-derived transcript expression values for each regulator and each target. The degree to which changes in expression of regulon components are consistent with the reported mode of action for each regulator-target pair (activator or silencer; dual regulators were ignored) was calculated by adjusting the sign of the log_2_-ratios (positive if the regulator’s mode of activity matches the expression change of the target, otherwise negative). The mean of directional log_2_-ratios was calculated to then evaluate the coherence of the regulator’s expression change with the concerted expression change across the regulator’s entire known regulon, here referred to as the “concerted log_2_-fold change in expression of regulon”. This value can then be compared to the log_2_-fold mutant versus wild-type expression change of the regulator for each regulon to assess whether there is evidence for a coherent increase or decrease in regulatory activity across a given regulon.

### Motility assays

Tryptone motility plates were made with 10 g/L of tryptone, 5 g/L of NaCl, and 3 g/L bacteriological agar, similar to those used in previously described motility assays^70^. MOPS-RDM and MOPS-Minimal media were also prepared and turned into motility plates by adding 3 g/L bacteriological agar and otherwise following the recipes listed above. Plates were poured evenly by serological pipette (20 mL/plate) and left on bench to dry overnight and then stored at 4C the next day. Plates were not used if more than 2 months had passed since pouring them.

Cryogenically preserved cells were streaked out on standard LB-agar plates, and then isolated colonies were grown in media of the same type for 12 hours in the case of LB/Tryptone broth and MOPS-RDM and 24 hours in the case of MOPS-Minimal. At the end of these growth periods, 1 mL of each culture was pelleted in a microcentrifuge for 3 minutes at 16,100xg at 4C. The supernatant was discarded, and the pellets were gently resuspended in 100 uL of sterile PBS pH 7.4. OD600 measurements were taken, and then additional PBS volumes were added to each resuspended pellet to normalize all samples to 80% of the OD600 of the lowest sample. Any condensate on the lid of the plates was wiped off using a sterile replica plating velvet. A small filter disk soaked with 10 uL of 10mM aspartic acid (as a chemoattractant) was then added to the center of MOPS-Minimal plates. 1 uL of each OD-normalized sample was then spotted onto all motility plates. We note to take care that the pipet tip should almost touch – but not break the surface tension – of the motility agar before dispensing the sample. After the plates were spotted, the plates were left facing up (media on bottom) and carefully parafilmed. The plates were then collectively placed into a Ziploc plastic bag containing some damp paper towels to prevent drying of plates. The bagged plates were then transferred to the 37C incubator for at least 12 hours before imaging at the first timepoint. After collecting the first timepoint, the plates were flipped upside-down (media on top) and were subsequently imaged every ∼2 hours.

To normalize the brightness of the motility plate images, the median brightness of each image was found using `imagemagick v7.1.04`^71^ `convert` with the `-colorspace gray` argument. The median of median brightnesses across all images was then calculated, and each image was adjusted in brightness to the value of the median of image-wise median brightness values using the `imagemagick convert -evaluate Multiply` argument.

## RESULTS

### Dam sites are statistically depleted in extended protein occupancy domains while controlling for AT content

We previously found, in wild-type *E. coli* K-12 MG1655, that there is a statistically significant underrepresentation of Dam sites within regions of the genome covered by EPODs relative to non-EPOD regions^46^. To control for the difference in AT% between EPODs and non-EPOD regions and address the possibility that the depletion of Dam sites in EPODs is caused by the AT-richness of EPODs, we compared the frequency of Dam sites within EPODs to the frequency of Dam sites at other loci of the same length as each EPOD (“shuffled” EPODs). We assigned each wild-type EPOD (“genomic” EPODs) to 10 equally populated bins defined by AT%, assigned 1000 permutations of shuffled EPODs to these AT% bins, and we performed Poisson regression analysis with terms for “genomic” (real) versus “shuffled” EPODs as well as AT% bin membership. Our results show that “genomic” EPODs contain significantly fewer Dam sites than “shuffled” EPODs while incorporating for the AT% bin membership term, and this is true both when shuffled EPODs are restricted from being shuffled to the positions originally occupied by genomic EPODs (−0.069 regression coefficient estimate for “genomic” EPODs, −0.120 to −0.017 95% confidence interval, 0.0104 p-value) and when shuffled EPODs are permitted to overlap genomic EPOD positions (−0.110 coefficient estimate for “genomic” EPODs, −0.170 to −0.064 95% confidence interval, 1.6 x 10^-5^ p-value). We performed this same Poisson regression analysis for Dcm sites and found no sign of anticorrelation for when we do not permit “shuffled” EPODs to overlap “genomic” EPODs (−0.011 coefficient estimate for “genomic” EPODs, −0.072 to 0.050 95% confidence interval, 0.734 p-value) and statistically significant anticorrelation – but much weaker than the anticorrelation for Dam sites – when we do allow “shuffled” EPODs to overlap “genomic” EPODs (−0.065 coefficient estimate, −0.130 to −0.004 95% confidence interval, 0.038 p-value). These results show that even after controlling for AT% the “genomic” EPODs have significantly fewer Dam sites than expected by random chance, thus reinforcing the basis of our hypothesis that there may be an association between EPOD formation – and more generally, protein occupancy – and DNA methylation (i.e., that the presence of Dam methylation might inhibit the NAP binding that gives rise to EPODs).

### Loss of DNA methylation minimally alters protein occupancy on the *E. coli* K-12 MG1655 genome

To characterize a set of protein binding events that may be dependent on DNA methylation state, we utilized the IPOD-HR methodology to profile changes in the global protein occupancy of the *E. coli* K-12 MG1655 chromosome when either or both of the genes encoding the two primary methyltransferases, *dam* and *dcm*, are deleted. IPOD-HR has been previously described and applied to characterize global protein occupancy changes in MG1655 NAP deletion mutants^46,47^. The IPOD-HR methodology involves crosslinking and Illumina sample preparation like other protein-DNA extraction and enrichment methodologies such as chromatin immunoprecipitation followed by sequencing (ChIP-seq)^72^. To produce a global profile of all protein occupancy across the genome, IPOD-HR utilizes physicochemical principles to enrich crosslinked protein-DNA complexes at an aqueous-organic interface during phenol/chloroform extraction. ChIP for RNA polymerase (RNAP-ChIP) is also performed on the same biological samples as used for IPOD-HR to remove the RNAP signal from the IPOD-HR occupancy profile. Producing RNAP-ChIP data allows us to isolate changes in occupancy of RNA polymerase (which we have observed to be correlated with gene expression changes) and the subtraction of the RNAP-ChIP signal from the IPOD-HR total protein signal highlights changes in transcription factor and NAP occupancy that might otherwise be obscured by replacement with RNAP^46^. We also performed RNAseq in parallel with IPOD-HR and RNAP-ChIP on *E. coli* MG1655 (WT) and our methyltransferase deletion mutants (*Δdam, Δdcm*, and *Δdam/Δdcm*) to characterize changes in gene expression that may result from changes in protein occupancy when DNA methylation is perturbed.

To explore the possibility of a global change in protein occupancy local to DNA methylation sites when the primary DNA methyltransferases are deleted, we first identified every potential Dam or Dcm site based on the appearance of their target motifs (5’-GATC-3’ or 5’-CCWGG-3’, respectively) in the *E. coli* K-12 MG1655 U00096.3 sequence. To capture the genomic context around each methylation site, we captured the 50 basepairs (bp) both upstream and downstream of each Dam or Dcm site, which generated 104 bp windows centered on each Dam site and 105 bp windows centered on each Dcm site. An association between methylation state and protein occupancy was then made by creating a distribution of the means of IPOD-HR or RNAP-ChIP occupancy scores within each methylation site window and comparing across genotypes (Figure 1A). We applied one-sample version of the Bayesian Estimation Supersedes the t-Test (BEST) analysis method to generate 95% credible intervals which for all conditions had a range of less than 0.1 and found that the distribution of occupancy scores is stable across genotypes. Overall, we observe no substantial differences in local IPOD-HR or RNAP-ChIP protein occupancy across all DNA methylation sites when *dam* and/or *dcm* are deleted.

**Figure 1:**
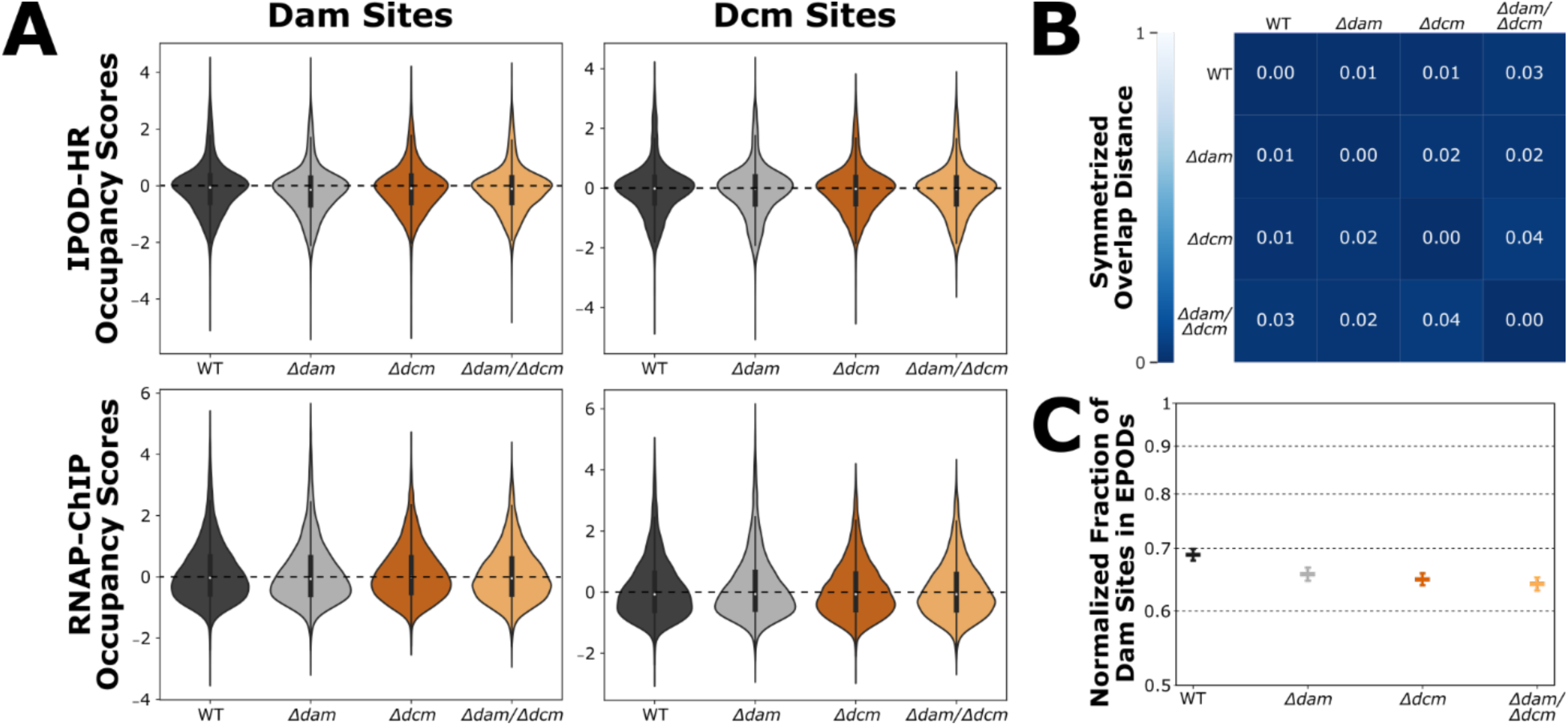
**(A)** Distribution of IPOD-HR and RNAP-ChIP occupancy scores in genomic windows centered on each Dam (104 bp window) or Dcm (105 bp window) target motif. **(B)** Symmetrized Overlap Distances calculated as described in Amemiya *et al.*, 2022^47^ to assess similarity in global EPOD composition between strains. A value of 0 indicates that all EPODs between the two strains overlap. **(C)** Normalized fraction of total Dam sites found within EPODs for each strain. The fraction of Dam sites within EPODs was normalized by the fraction of total genomic basepairs covered by EPODs, and 95% confidence intervals were calculated through jackknife resampling of the EPOD genomic positions. The y-axis is log- scaled.

The statistical under-representation of Dam sites in EPODs also motivated us to explore how EPOD locations change relative to DNA methylation sites when *dam* and/or *dcm* are deleted. To characterize the set of EPOD locations for each genotype, EPOD calling was performed on IPOD-HR data as previously reported with some minor modifications (as noted in Methods)^46^. Symmetrized Overlap Distances (SODs) were calculated as previously described in Amemiya *et al*., 2022^47^ to analyze similarity in the set of EPOD locations between genotypes (Figure 1B). A SOD score of “0” represents perfect overlap of EPOD locations between sets and a score of “1” represents zero overlap in EPOD locations between sets. Given that the SOD scores between any compared genotypes were 0.04 or lower, we observe negligible changes in the set of EPOD locations when *dam* and/or *dcm* are deleted. We additionally found no substantial change between genotypes in the fraction of Dam or Dcm target sites within EPODs (Figure 1C), although a slight decrease in the fraction of Dam sites in EPODs is apparent. These results indicate that there are few to no changes in large-scale protein occupancy features, such as EPOD formation, when DNA methylation is perturbed through methyltransferase deletion.

To evaluate whether the loss of methylation signals might still alter gene expression within EPODs when compared to non-EPOD regions of the genome, we performed RNA-seq experiments on all four strains included in our study and calculated the median mutant versus wild-type log_2_FC of transcripts within EPODs and subtracted the median mutant versus wild-type log_2_FC of transcripts outside of EPODs. We found a small but significant decrease in median expression of transcripts within EPODs versus outside of EPODs when *dam* and both *dam* and *dcm* deleted, but there is no difference in median expression between within and outside of EPODs when just *dcm* is deleted (median difference of mutant versus wild-type log_2_FC of transcripts within EPODs minus outside of EPODs and Wilcoxon two-sided rank-sum p-value: 0.06 and 1.1 x 10^-5^ for *Δdam*, 0.00 and 0.95 for *Δdcm*, 0.15 and 3.2 x 10^-8^ for *Δdam/Δdcm*). Thus, there is a small relative decrease in transcription inside vs. outside of EPODs when *dam* is deleted, although especially considering the absence of systematic occupancy changes, the mechanism and biological significance of these changes remains unclear. Additional analysis of our RNA-seq results is provided below.

### Protein occupancy signal in *dam* deletion mutants is decreased at dense clusters of Dam target sites

Considering that multiple DNA methylation events in close genomic proximity could induce a greater degree of DNA curvature^73^, we hypothesized that genomic regions with dense clusters of methylation sites may experience more pronounced protein occupancy changes when DNA methylation is perturbed. In addition, it has been shown that certain DNA-binding proteins such as SeqA preferentially bind to regions with multiple proximal Dam sites^21,74^. These considerations led us to consider how the change in IPOD-HR occupancy differences between methyltransferase deletion mutants and the wild type might vary as a function of local methylation site density. Here we observe a negative correlation between mutant-specific IPOD-HR signal and Dam Site Density when *dam* is deleted. In other words, we find less total protein occupancy at dense clusters of Dam sites when *dam* is deleted, whereas locations with lower Dam site densities are unaffected, as are Dcm sites.

Our analysis defined twelve unique loci containing a Dam site annotated with a Dam Site Density of 6 as well as three unique loci with a Dam site annotated with a Dam Site Density of 7; we refer to these regions collectively as “high-density Dam site clusters”. Each of these high-density Dam site clusters appear within ORFs and are thus absent from promoters or intergenic regions. Despite the *Δdam*-associated increase in RNA polymerase at most of these high-density Dam site clusters, there is only one locus which presents with a significant change in proximal gene expression, which is the *oriC-*adjacent gene *mnmG* (Rockhopper q-values for transcript count of *mnmG*: ∼.0017 for *Δdam* vs. WT, ∼1.0 for *Δdcm* vs. WT, ∼.00036 for *Δdam/Δdcm* vs. WT). Overall, it appears that the *Δdam*-dependent change in RNAP and total protein occupancy at high density Dam site clusters are not impacting expression of known local transcripts (see GEO dataset GSE279866).

One locus of interest featuring a cluster of seven Dam sites is at the terminal end of the *selB* coding region (Figure 2BC). Here we observe *Δdam*-dependent peaks in RNAP occupancy both at the Dam site cluster as well as at the promoter immediately upstream of *selB*. However, transcript levels of *selB* do not change substantially in *Δdam* genotypes (Rockhopper mutants versus wild-type log_2_FC / q-values of *selB*: −0.21 / 0.69 for *Δdam* vs. WT, 0.072 / 0.27 for *Δdcm* vs. WT, −0.16 / 3.6 x 10^-4^ for *Δdam/Δdcm* vs. WT). The gene with a promoter immediately downstream of the *Δdam*-dependent RNAP-ChIP peak at the *selB* Dam site cluster, *yiaY*, is not transcribed in any of our genotypes (see GEO dataset GSE279866). There is a decrease in IPOD-HR signal associated with the increase in RNAP-ChIP signal at the *selB* Dam site cluster in *Δdam* strains, but we cannot find any reports on what protein may bind to this region.

**Figure 2:**
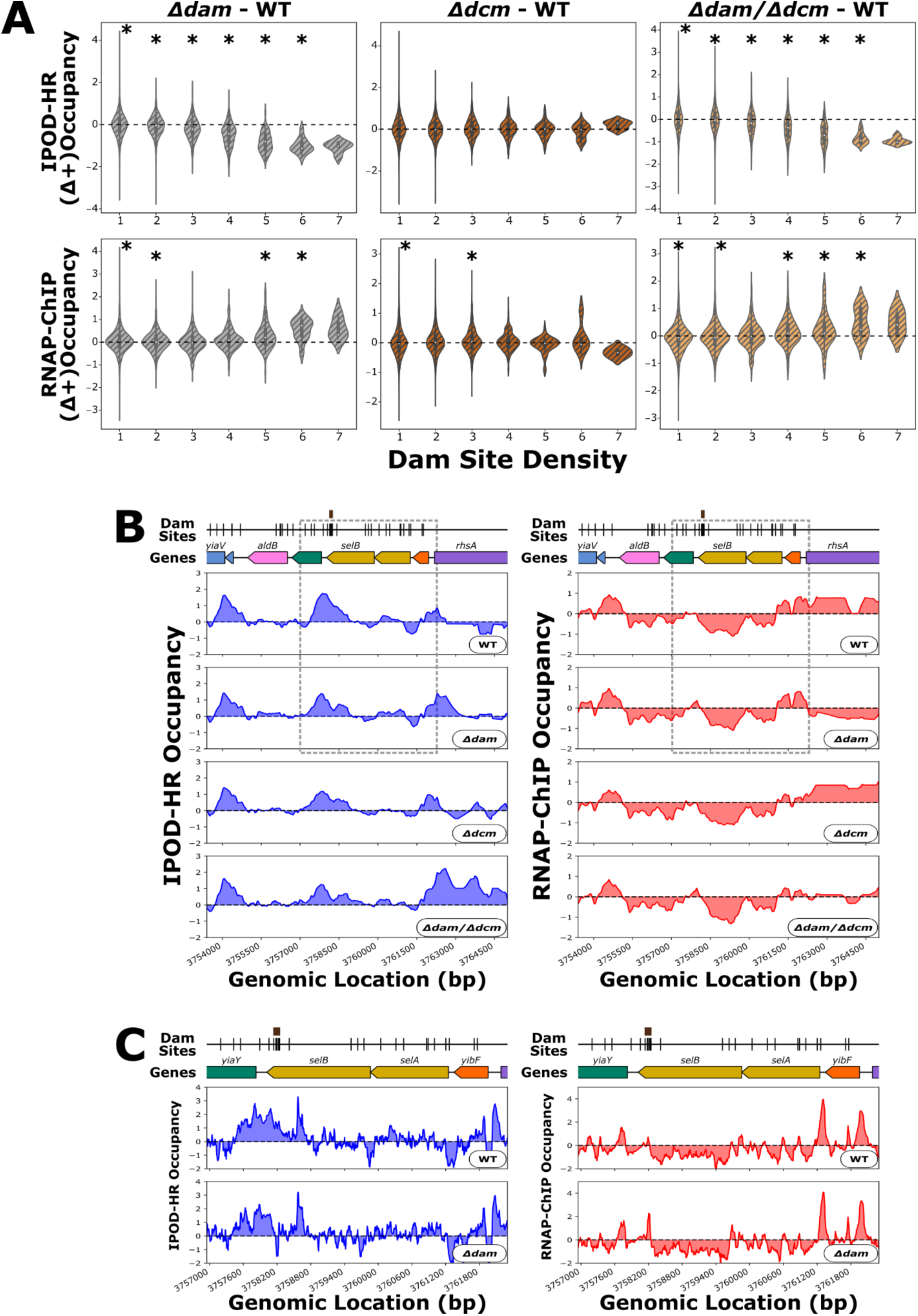
**(A)** Distribution of changes in mean IPOD-HR or RNAP-ChIP occupancy scores in 104 bp windows centered on each Dam target motif; positive scores indicate higher occupancy in the indicated mutant relative to WT. “Site Density” on the x-axis refers to the number of Dam sites within each window. Asterisks represent p-values of < 0.01 by Wilcoxon signed rank test with Bonferroni correction. At the higher site densities (6 and 7 site density) there is a lack of statistical power due to a small number of loci with such high methylation site densities. **(B)** Genomic context of *selB* showing 512 bp rolling mean of IPOD-HR (blue occupancy trace) or RNAP-ChIP (red occupancy trace) robust z-scores. Brown boxes above markers on the “Dam Sites” tracks indicate “7 Dam Site Density” clusters of interest. Genes are differentially colored based on their membership to functional gene clusters. The dashed box designates the locus which is shown in panel C. **(C)** Zoomed in view of *selB* (corresponding to the boxed region of panel **B**) showing IPOD-HR (blue occupancy trace) or RNAP-ChIP (red occupancy trace) robust z-scores.

We also identified a 7 Dam site cluster at the terminal end of *prpE* (Figure 3AB), which is comparable to the *selB* case in that it features an increase in RNAP-ChIP signal and a decrease in IPOD-HR signal in our *Δdam* genotypes. In contrast to the *selB* case, for *prpE* the *Δdam*-dependent peak in RNAP-ChIP appears ∼200 basepairs downstream of the Dam site cluster, and so the RNAP-ChIP peak is proximal to the promoter for *codB*, which shows modest but not statistically significant increases in transcript levels in all of our methyltransferase deletion strains (Rockhopper mutants versus wild-type log_2_FC / q-values of *codB*: 0.30 / 1.0 for *Δdam* vs. WT, 0.45 / 1.0 for *Δdcm* vs. WT, 0.31 / 0.11 for *Δdam/Δdcm* vs. WT). Given that there is a similar magnitude of upregulation of *codB* in *Δdcm* as compared to *Δdam*, the upregulation of *codB* may not actually be associated with the *Δdam*-dependent increase in RNAP-ChIP, but rather is likely statistical noise. *prpE* is not actively transcribed in any of our strains (see Supplementary Data).

**Figure 3:**
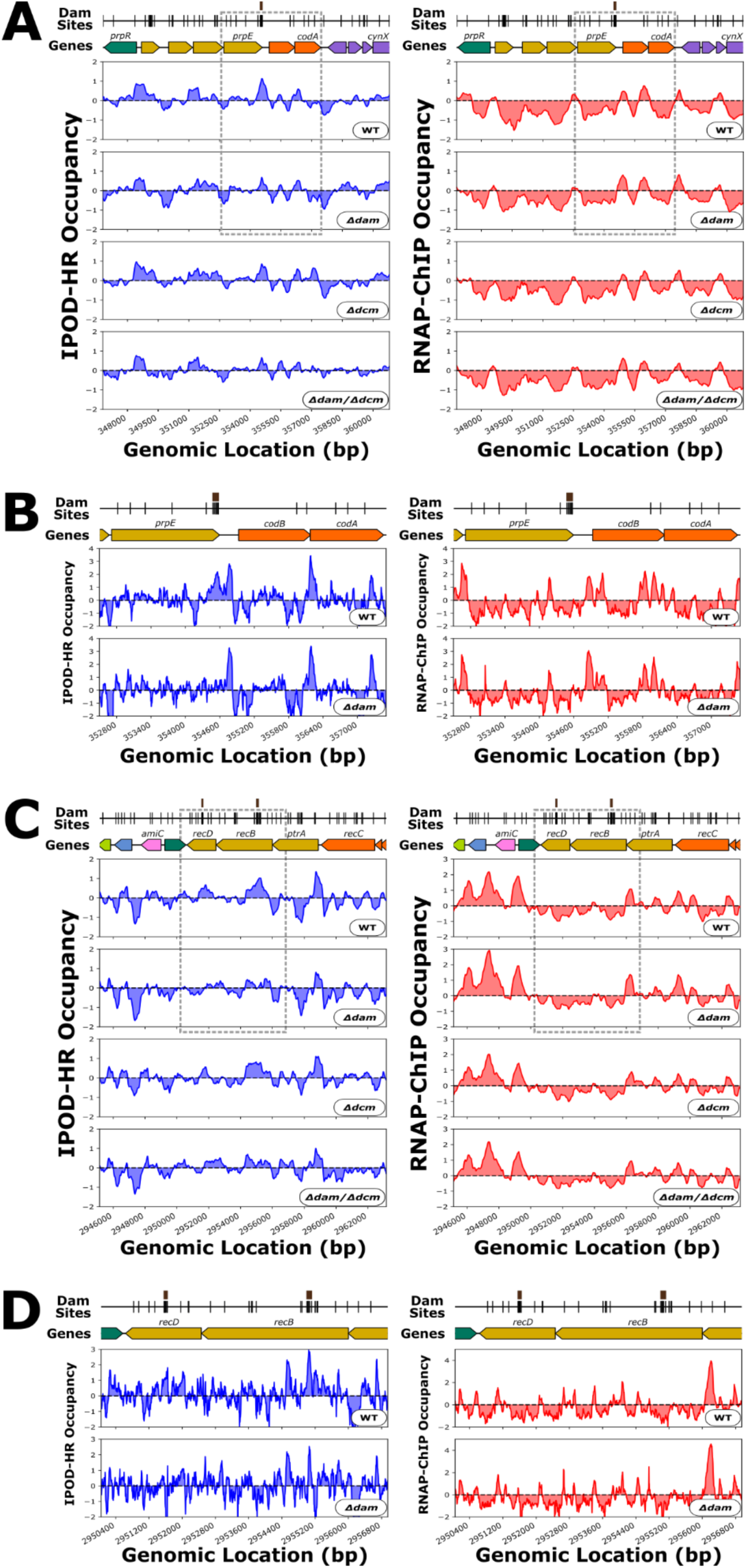
**(A)** Genomic context of *prpE* showing 512 bp rolling mean of IPOD-HR (blue occupancy trace) or RNAP- ChIP (red occupancy trace) robust z-scores. Brown boxes above markers on the “Dam Sites” tracks indicate “7 Dam Site Density” clusters of interest. Genes are differentially colored based on their membership to functional gene clusters. The dashed box designates the locus which is shown in panel **B**. **(B)** IPOD-HR (blue occupancy trace) or RNAP-ChIP (red occupancy trace) robust z-scores in the immediate vicinity of *prpE*. **(C)** As in panel **A**, showing the genomic context around *recBD*. **(D)** As in panel **B**, for the boxed region indicated in panel **D**.

There are two proximal 7 Dam site clusters within the *recB* and *recD* coding regions (Figure 3CD). We find this notable because of the involvement of *dam*, *recB*, and *recD* in *E. coli* DNA mismatch repair, and the fact that previous attempts to delete both *dam* and *recB* found such a strain to be inviable^75^. Despite an increase in the RNAP-ChIP signal at the *recBD* promoter in our *Δdam* genotype, the transcript levels of *recB* in fact show a small decrease in our *dam* mutant strains (Rockhopper mutants versus wild-type log_2_FC / q-values of *recB*: −0.043 / 0.65 for *Δdam* vs. WT, 0.082 / 0.14 for *Δdcm* vs. WT, −0.087 / 7.7 x 10^-4^ for *Δdam/Δdcm* vs. WT) and *recD* (Rockhopper mutants versus wild-type log_2_FC / q-values of *recD*: −0.31 / 0.0052 for *Δdam* vs. WT, −0.057 / 0.075 for *Δdcm* vs. WT, −0.31 / 7.1 x 10^-5^ for *Δdam/Δdcm* vs. WT). The *recD* Dam site cluster is in the middle of the gene body while the 7 Dam site cluster in *recB* is a few hundred basepairs downstream of a putative *recB* promoter (defined by the presence of an RNA polymerase ChIP peak), and the IPOD-HR occupancy at both sites is substantially decreased in our *Δdam* genotype.

Rifampicin – at the concentration added to our cells used for RNAP-ChIP – prevents promoter clearance, which leads to a build-up of RNAP at active promoters^76,77^. We thus hypothesized that the *Δdam*-dependent RNAP-ChIP peak observed at these Dam site clusters may result from RNAP that has been directly recruited for transcription – perhaps of a small RNA. However, there does not appear to be any increase in RNA-seq reads local to the RNAP-ChIP peaks at these Dam site clusters in *Δdam* strains (Supplementary Figure 1). We also considered that these RNAP-ChIP peaks may result from RNAP being stalled, possibly at a Dam methylation-directed repair site due to the increase in DNA damage and upregulation of SOS response in *Δdam* strains^78–80^. To investigate the possibility of DNA damage, we produced heatmaps of the normalized frequency of read ends at some of the 7 density Dam site clusters, but none of the heatmaps show a *Δdam*-dependent pattern in read end accumulation local to the RNAP-ChIP peaks (Supplementary Figure 2), indicating no clear signature of increased strand breaks near the Dam site cluster (although any such accumulation may well have been obscured anyway by the fragmentation steps inherent to our purification protocols). Taken together, these results indicate that RNAP may be directly recruited to – but not actively transcribing – the high-density Dam site clusters in *Δdam* strains, or RNAP may be stalled but not because of DNA strand breaks; additional investigation would be required to distinguish between these possibilities.

### Methyltransferase deletion globally perturbs expression of multiple large regulons

While global protein occupancy is generally stable across most methylation sites when DNA methylation is perturbed, deletion of *dam* and/or *dcm* has been associated with some changes in gene expression^21,22,38,81^. We produced RNAseq data which was analyzed through the Rockhopper analysis suite^64–66^ and found gene expression changes across many operons in the *Δdam* and *Δdam/Δdcm* genotypes, while the *Δdcm* strain showed relatively fewer significant changes in gene expression (Figure 4). In the *Δdam* single mutant, the most positively expressed genes relative to wild-type are associated with DNA damage response, and the most down-regulated genes are primarily members of the *gatYZABCDR* operon which is involved in galactitol catabolism^82^ (Table 1). Both the most up-expressed and most down-expressed genes in *Δdcm* encode gene products for transmembrane transport. Aside from the genes already represented in the single deletion mutants, the double deletion mutant *Δdam/Δdcm* is characterized by upregulation of maltooligosaccharide catabolism proteins encoded by *malP* and *malQ*^83^ as well as downregulation of isoleucine and valine biosynthesis through *ivbL*^84^.

**Figure 4:**
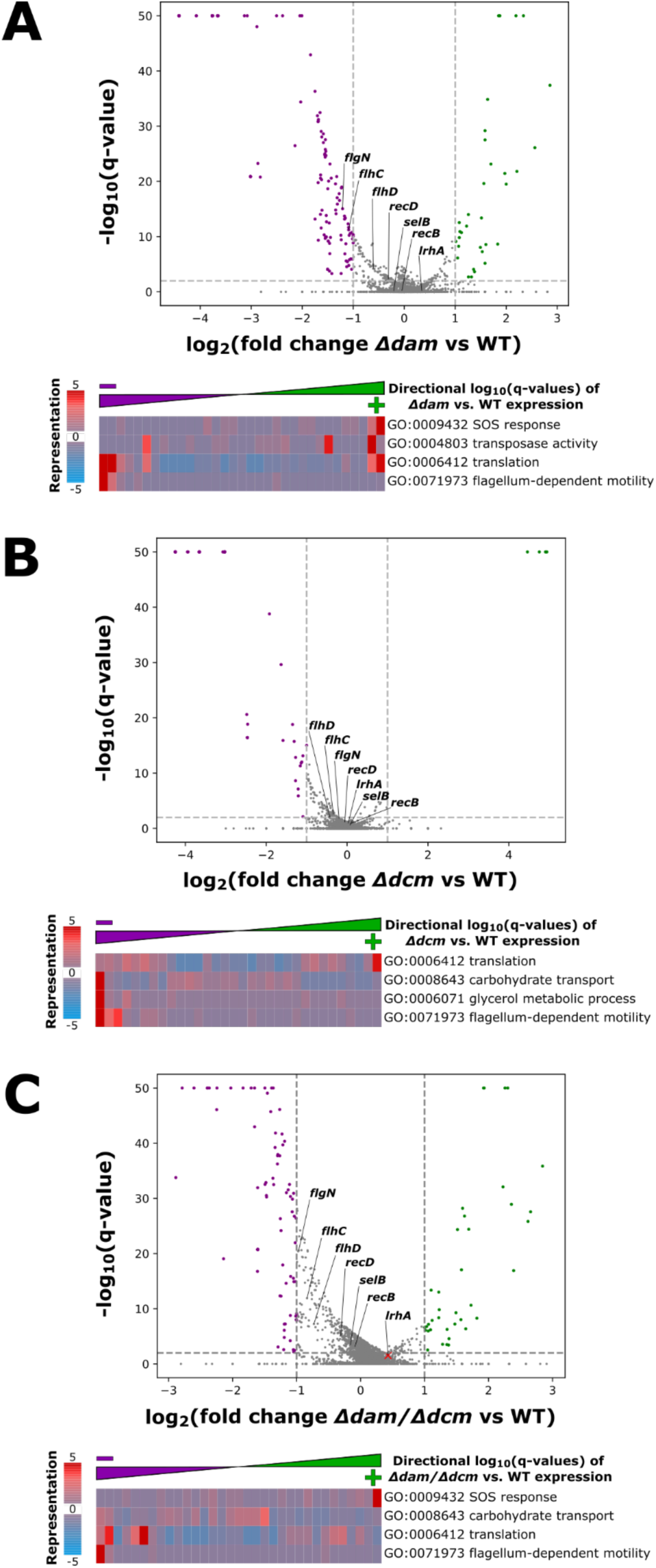
**(A)** *Δdam*, **(B)** *Δdcm,* and **(C)** *Δdam/Δdcm* versus wild-type change in expression of genes. Purple genes are reduced in transcript abundance – while green genes are increased in abundance – in the mutant relative to wild-type. Some genes of interest are labeled by name. Below each volcano plot is shown gene set enrichment analysis for RNA-seq data across the indicated genotype (relative to wild type) for a selected subset of gene ontology terms. iPAGE reports the representation of directional log10(q-values) across the genes annotated with each Gene Ontology (GO) term – thus, a redder bin indicates an over-representation of genes from the specified GO-term (row) at that expression change bracket (column).

**Table 1:**
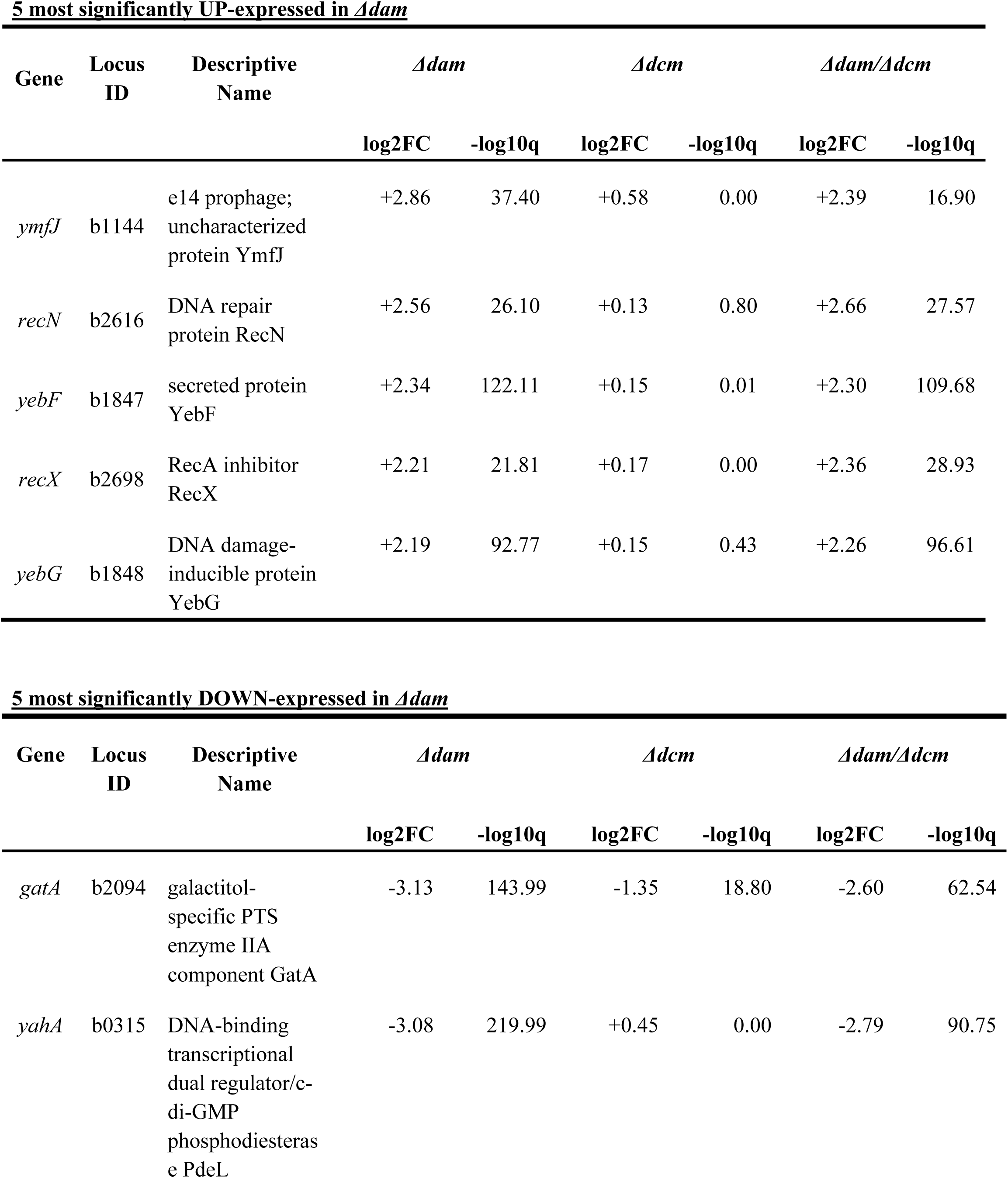
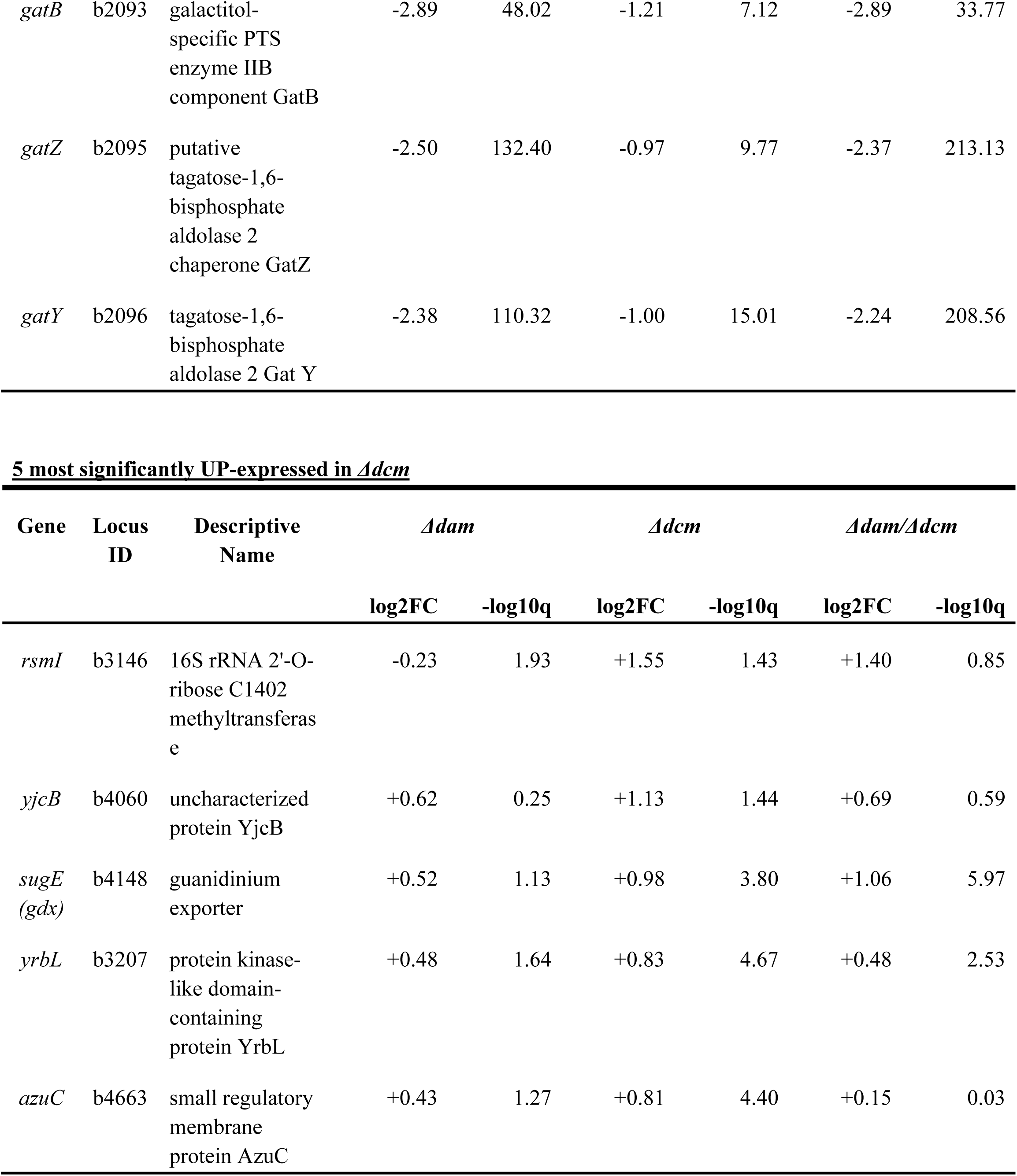
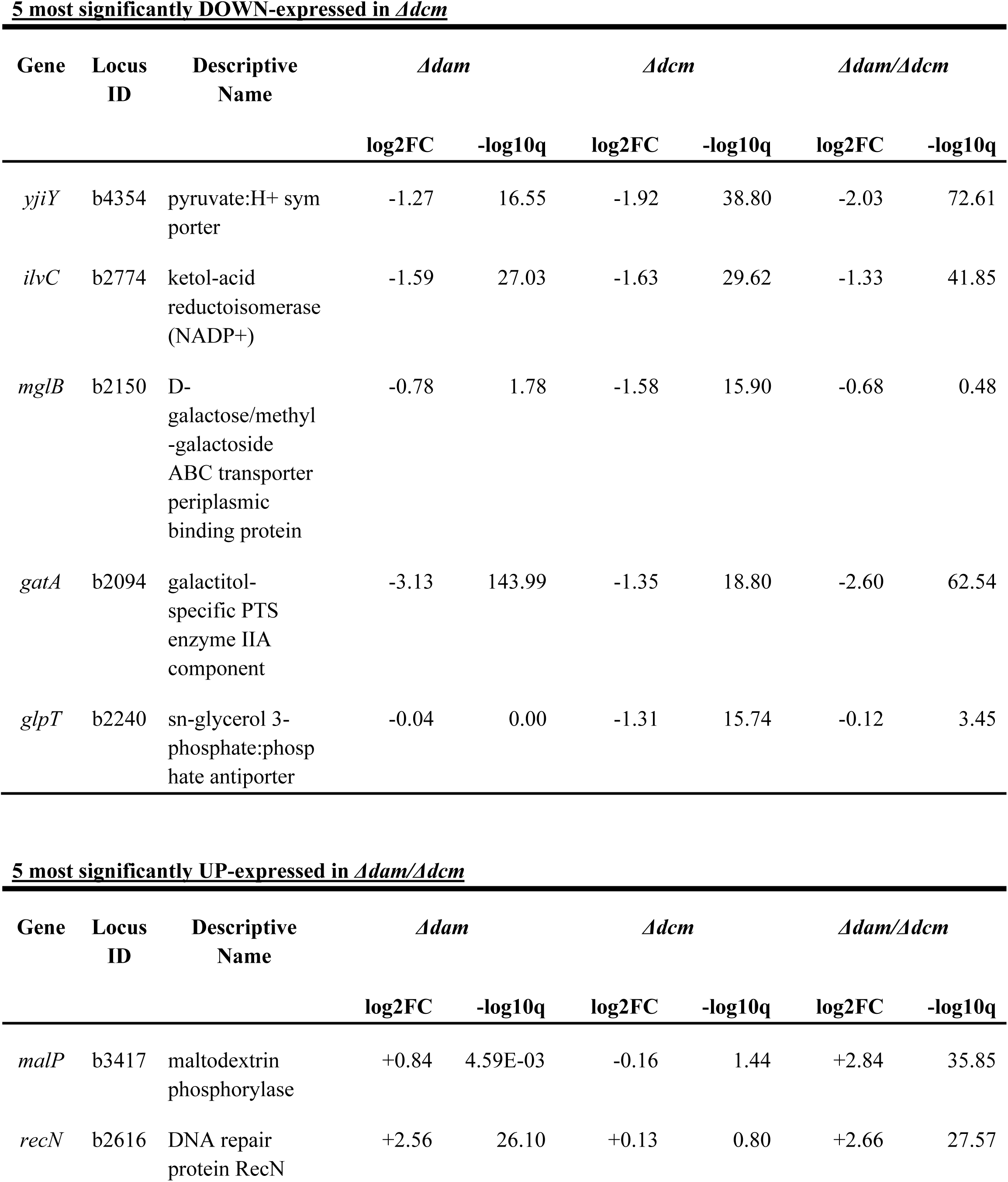
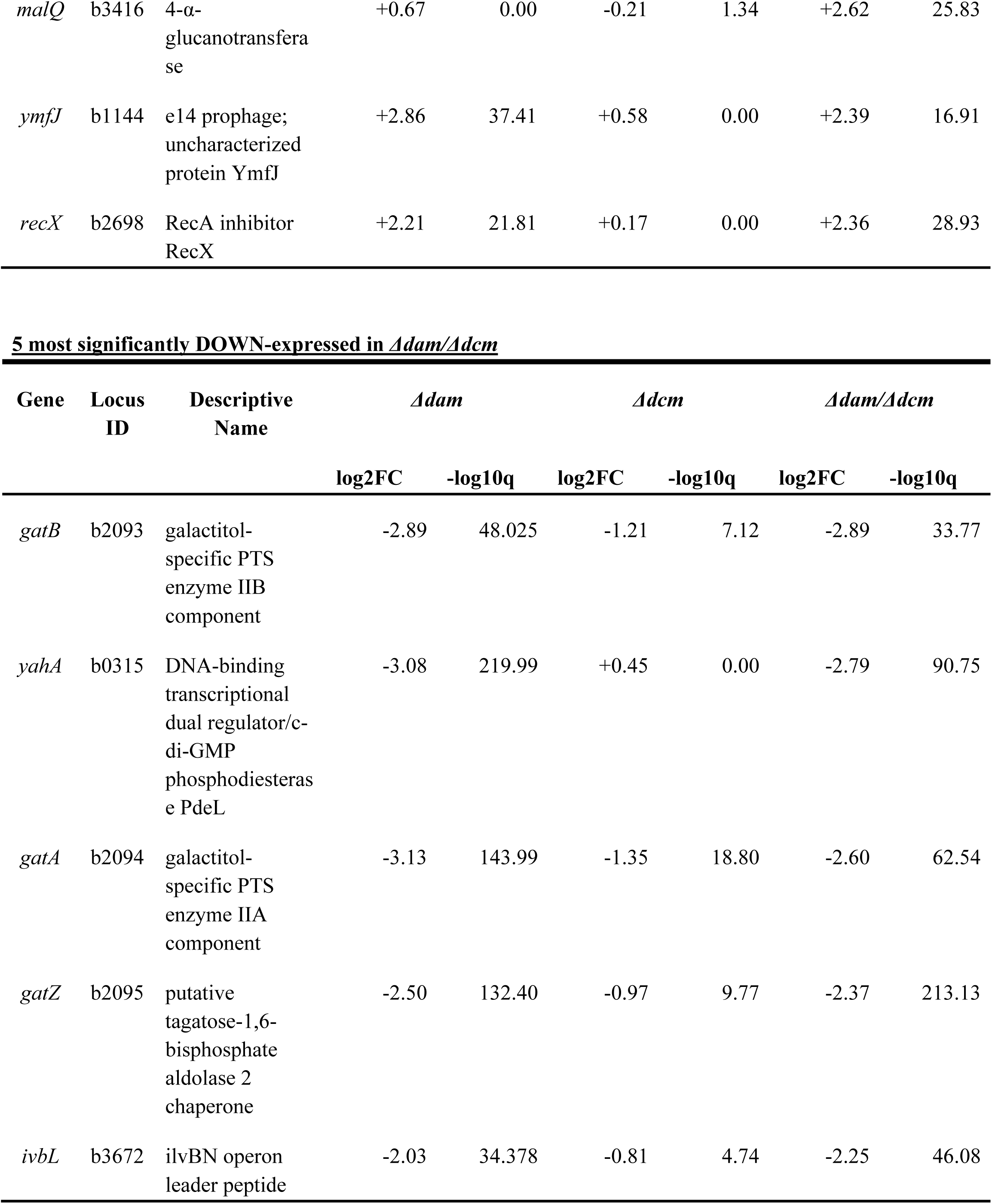
Transcripts showing the largest changes in abundance in each mutant genotype relative to wild-type, as calculated using Rockhopper.

We then performed gene set enrichment analysis on our RNAseq data which further supports gene expression changes across multiple gene ontology (GO) categories in methyltransferase deletion mutants (Figure 4). We found a decrease in expression of several genes associated with flagellum-dependent motility which was consistent in all methyltransferase mutants relative to wild-type. Previous findings that activation of SOS response, which is associated with DNA damage repair, occurs upon *dam* deletion were also reproduced here, likely caused by interference with hemimethylation-dependent mismatch repair and/or perturbation of normal replication initiation^21,85^. We also detected differential expression of gene products involved in maintaining transposons across all methyltransferase mutants, which supports previous findings that *dam* methylation impacts transposase expression and transposition activity^86,87^. Genes relating to translation and amino acid biosynthesis have also been reported to significantly change in expression in *Δdam* strains^20,80^, and our methyltransferase mutants all show substantial expression changes in translation and amino acid biosynthesis (Supplementary Figure 3), but the mechanistic basis underlying the relationship between transcription of translation-related and DNA methylation has not been elucidated.

### Previously identified instances of local methylation-sensitive regulation are recapitulated in our data

To systematically investigate the concordance of our data with previous reports of correlations between methylation and protein occupancy in *E. coli* strains^18,19,22,88–91^, we first reviewed the available literature to compile a list of candidate genes which had previously been studied in the context of potential contributions of Dam methylation to their cis-regulatory logic, requiring that those genes had (a) been investigated for change in transcript level based on methylation state of immediately upstream Dam or Dcm sites, or (b) had been reported as having persistently unmethylated immediately upstream Dam or Dcm motifs (Table 2). We included data arising both from genetic deletions of *dam* or *dcm*, and from 5-azacytidine (5-aza) treatments, which blocks the addition of methyl groups to cytosine by Dcm^23,92^.

**Table 2:**
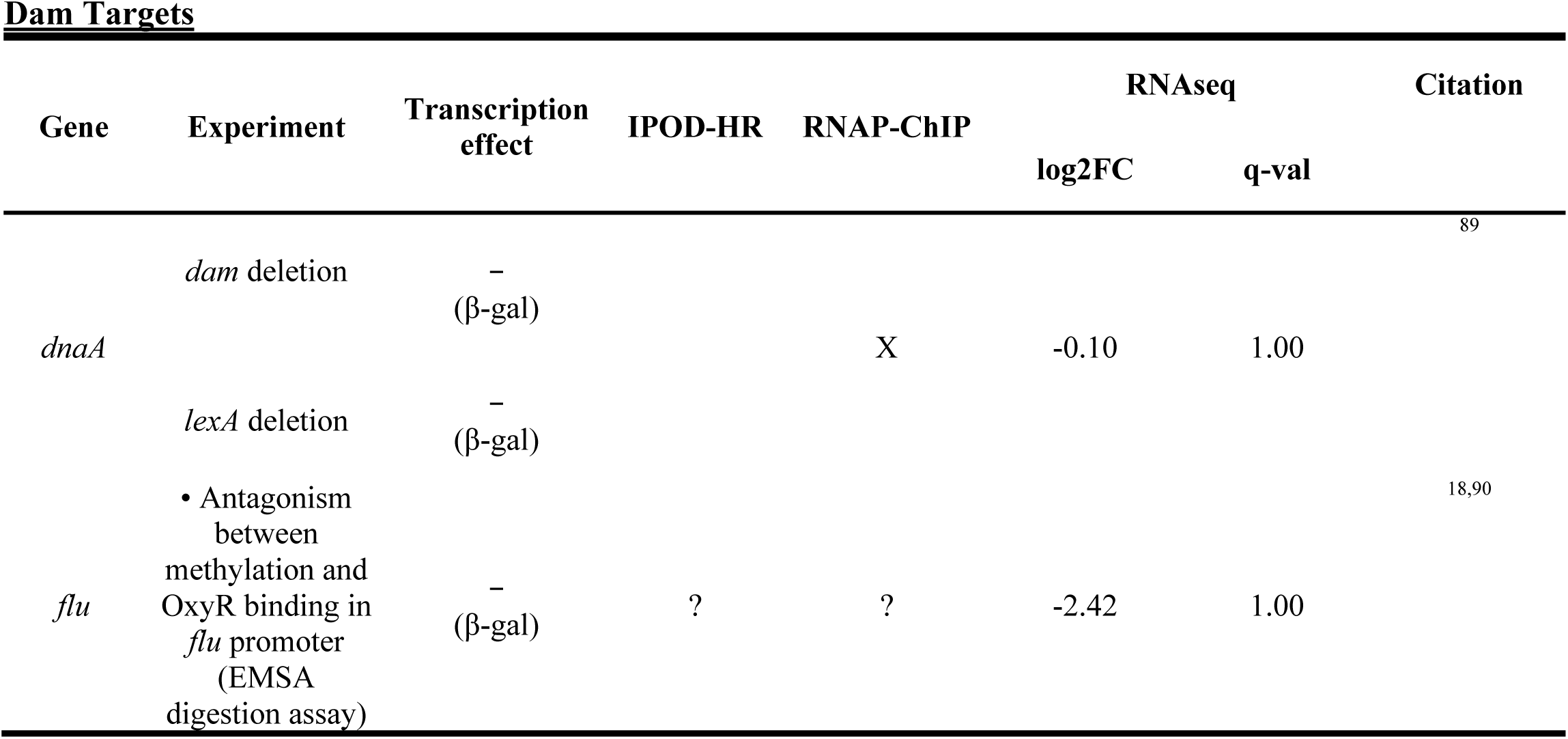
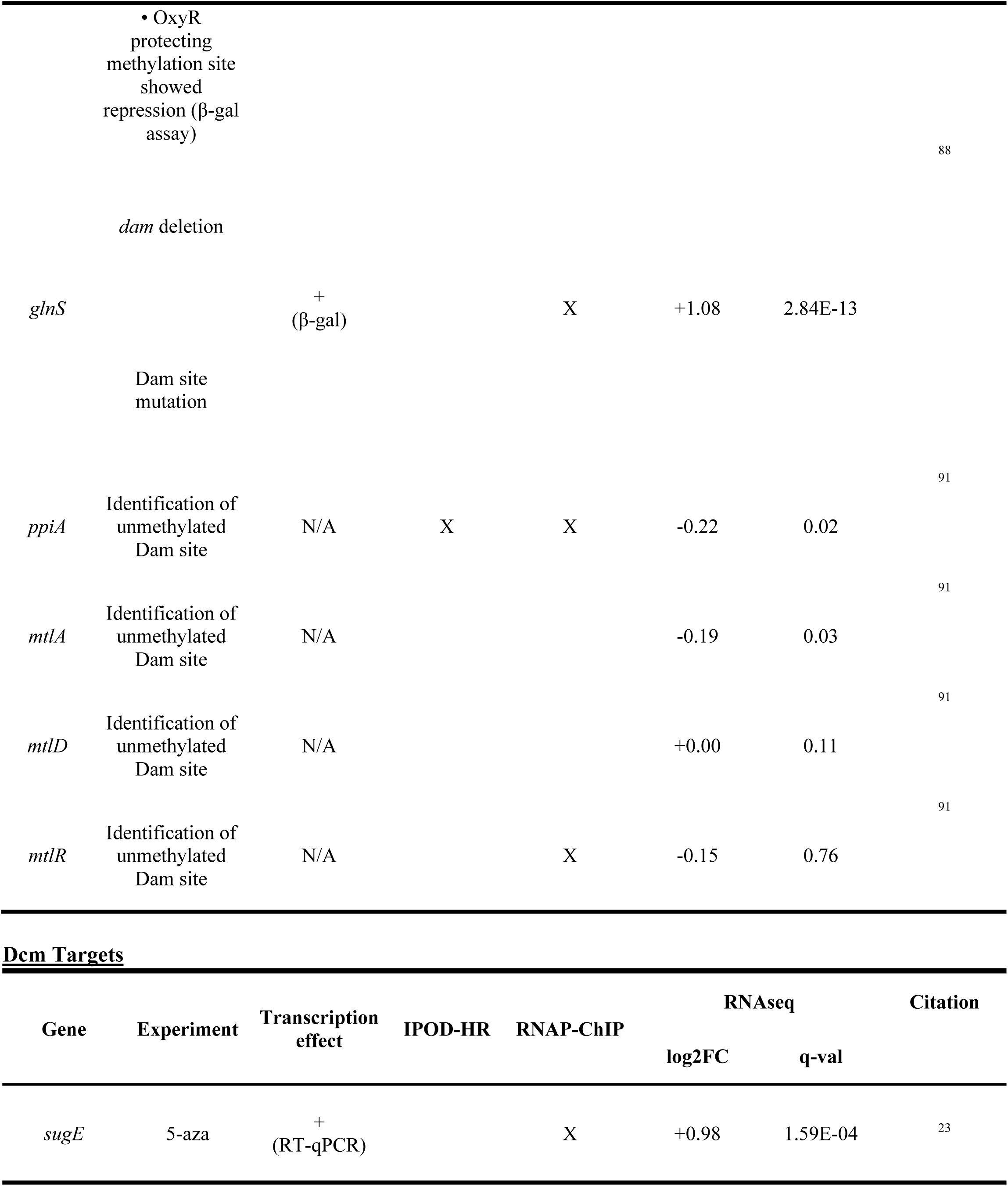
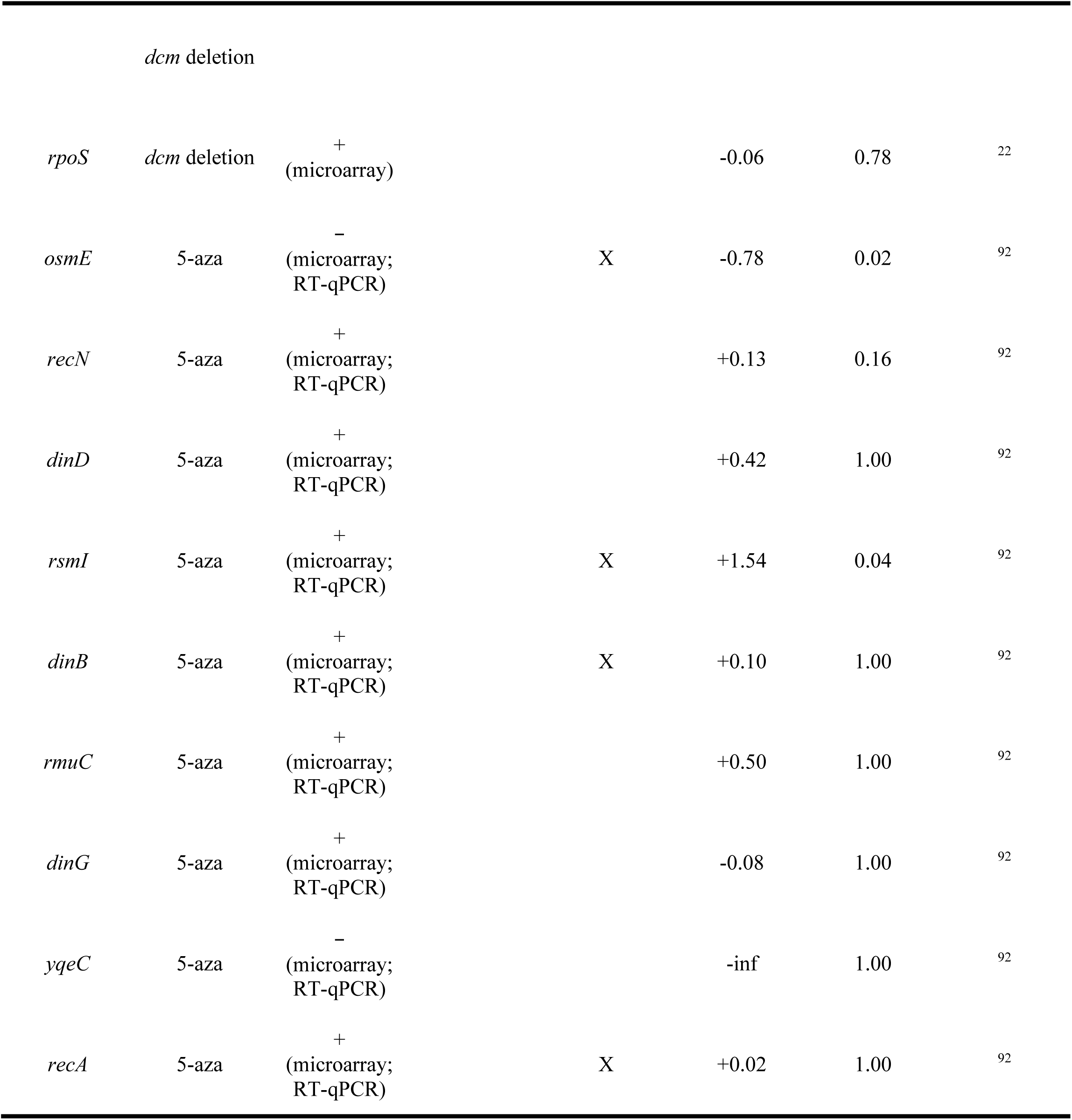
Summary of candidate genes for regulation by local Dam and Dcm methylation based on available literature, compared with our IPOD-HR, RNAP-ChIP, and RNAseq results. The “Experiment” column briefly describes the conditions in the citation that perturb methylation, and “Transcription effect” refers to the directionality of the change in expression (and methodology used to measure expression change) for the “Gene” when methylation is lost as reported in the citation. “X” indicates an observable change in occupancy pattern for the respective mutant relative to wild-type, and “?” indicates noisy signal at that locus which precludes interpretation. RNAseq results are also reported with respect to mutant versus wild-type.

Militello *et al*., 2014^23^ identified that in both *dcm* deletion strain and in WT cells subjected to 5-azacytidine treatment, *sugE* is derepressed – our results agree with this finding as despite the *Δdcm*-dependent decrease in RNAP-ChIP occupancy at the *sugE* promoter (Figure 5AB) there is still an increase in *sugE* transcript levels in our *Δdcm* genotypes (Figure 5C). Conversely, Militello *et al*., 2014^23^ reported that there are several Dcm sites in the *sugE* promoter and body but the genome of *E. coli* MG1655 does not contain any Dcm sites proximal to *sugE*, and so any relation between Dcm activity and *sugE* expression does not appear to result from occupancy changes dependent on local methylation state, but rather is likely an indirect regulatory effect.

**Figure 5:**
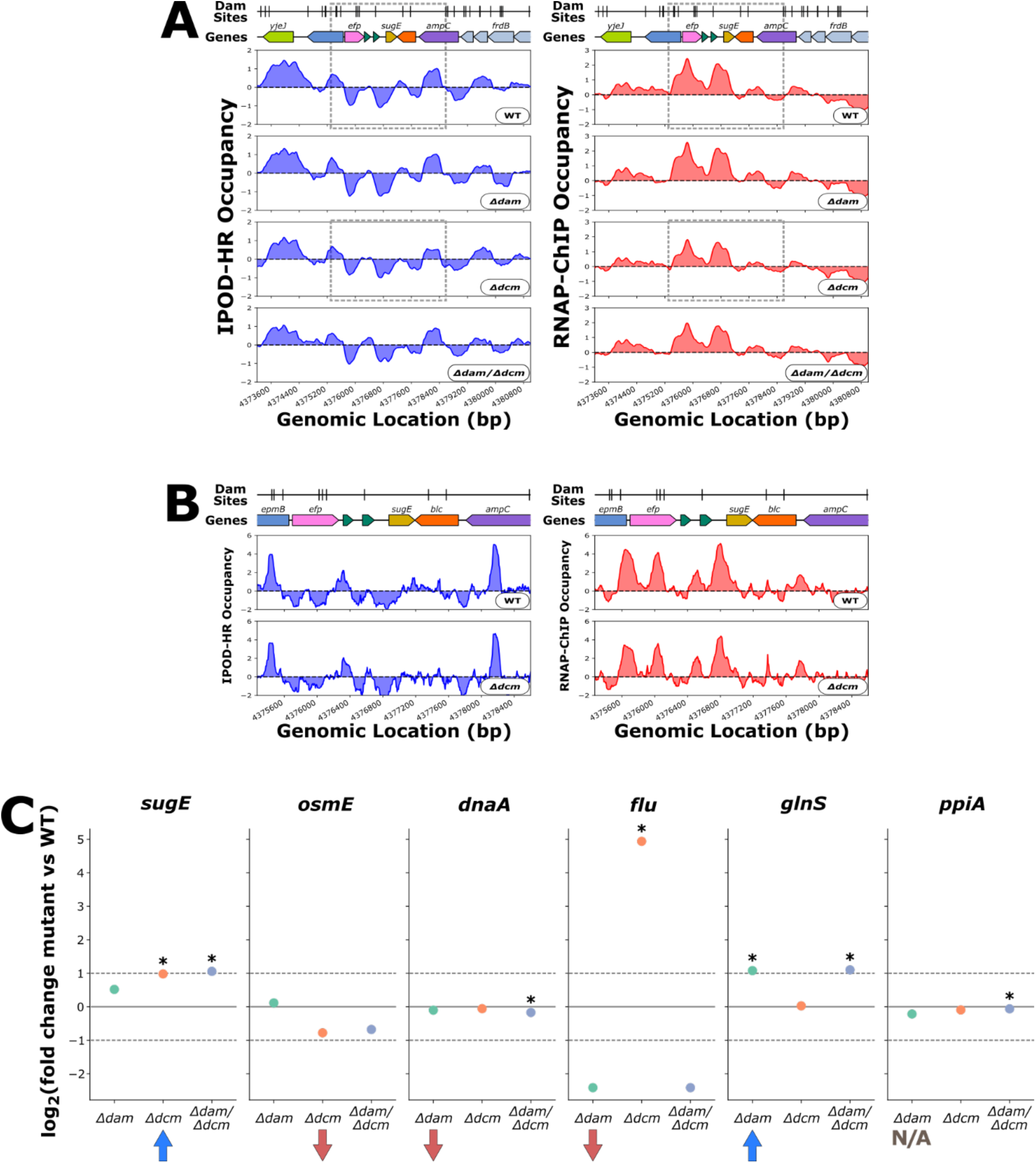
**(A)** Genomic context of *sugE* showing 512 bp rolling mean of IPOD-HR (blue occupancy trace) or RNAP-ChIP (red occupancy trace) robust z-scores. Genes are differentially colored based on their membership to functional gene clusters. The dashed box designates the locus which is shown in panel B. **(B)** Genomic locus of *sugE* showing IPOD-HR (blue occupancy trace) or RNAP-ChIP (red occupancy trace) robust z-scores. **(C)** Mutant versus wild-type change in expression where the red and blue arrows and gray “N/A” indicate the expected change in expression based on curated literature for *sugE*^23^, *osmE*^92^, *dnaA*^89^, *flu*^18,90^, *glnS*^88^, and *ppiA*^91^. Asterisks indicate statistical significance with a q-value less than 0.05 as calculated by Rockhopper.

In a similar study, Militello *et al*., 2016^92^ found that 5-azacytidine treatment increases transcript levels of *recN*, *dinD*, *dinG*, *rsmI*, *dinB*, *rmuC*, and *recA*, and decreases transcript levels *osmE* and *yqeC*. Our *dcm* deletion results are consistent with these findings (in terms of the sign of the log fold change upon *dcm* deletion) for all of the genes showing increased transcript levels in ^92^ except for *dinG,* which we find to drop in expression in our *Δdcm* strain, albeit not significantly (Table 2). For *osmE* and *yqeC*, our *Δdcm* strain shows repression in agreement with the 5-aza study, but *yqeC* is not expressed in any of our strains under our conditions. Therefore our *dcm* deletion genotype mostly recapitulates the expression changes observed from 5-aza treatment in Militello *et al*., 2016^92^ except for in the case of *dinG* where we report an opposing impact on gene expression.

Several similar datasets have been obtained to study specific instances of regulation of transcription by Dam. Braun and Wright 1986^89^ conducted an *in vivo* β-Galactosidase activity assay and S1 nuclease mapping in addition to *in vitro* transcription run-off experiments which all supported that loss of Dam methylation in the *dnaA* promoter leads to repression of *dnaA*. We find repression of *dnaA* in our *Δdam* genotypes, but we could not identify any protein occupancy change proximal to the *dnaA* promoter between our wild-type and *Δdam* strains in our IPOD-HR results, possibly due to the competition between DnaA and SeqA for binding to this region^93,94^ (as IPOD-HR would not distinguish between the two factors) or the fact that all of our experiments are ensemble averages over actively growing populations. Correnti *et al*., 2002^90^ and Wallecha *et al.*, 2002^18^ both investigated antagonism between OxyR and Dam methylation in the promoter of *flu* where loss of methylation led to OxyR binding which led to *flu* repression. Our data supports that *dam* deletion is associated with *flu* repression, but further assessment of OxyR-methylation antagonism is made difficult due to noisy IPOD-HR and RNAP-ChIP signal at the *flu* promoter which may be due to issues in data processing and quantitation as a result of repetitive sequence from the transposable element present in that region. Plumbridge and Söll 1987^88^ performed *in vivo* β-Galactosidase activity assays which showed that *dam* deletion as well as mutation of Dam sites in the promoter leads to derepression of *glnS*. Here we find, consistently, that *glnS* is more strongly expressed in our *Δdam* strains. It is, however, notable that there is a marginal increase in RNAP-ChIP occupancy at the *glnS* promoter in only the *Δdam* single deletion mutant and IPOD-HR occupancy at this locus is roughly static across our genotypes (Figure 6AB).

**Figure 6:**
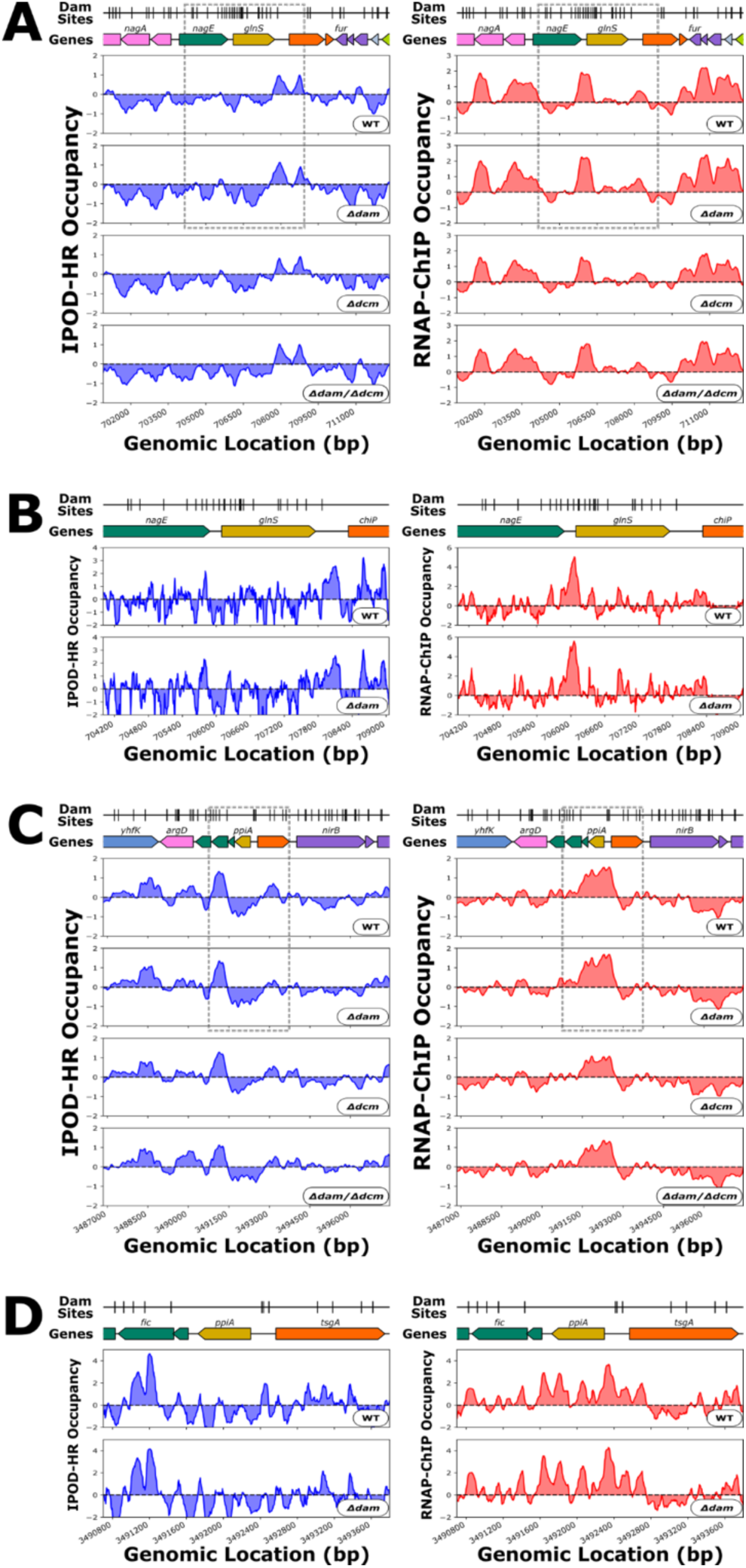
**(A)** Genomic context of *glnS* showing 512 bp rolling mean of IPOD-HR (blue occupancy trace) or RNAP-ChIP (red occupancy trace) robust z-scores. Genes are differentially colored based on their membership to functional gene clusters. The dashed box designates the locus which is shown in panel B. **(B)** Genomic locus of *glnS* showing IPOD-HR (blue occupancy trace) or RNAP-ChIP (red occupancy trace) robust z-scores. **(C)** Genomic context of *ppiA* showing 512 bp rolling mean of IPOD-HR (blue occupancy trace) or RNAP-ChIP (red occupancy trace) robust z-scores. Genes are differentially colored based on their membership to functional gene clusters. The dashed box designates the locus which is shown in panel D. **(D)** Genomic locus of *ppiA* showing IPOD-HR (blue occupancy trace) or RNAP-ChIP (red occupancy trace) robust z-scores.

Hale *et al*., 2004^91^ identified under various growth conditions the Dam sites that remain specifically unmethylated throughout the cell cycle. Of the genes reported to be proximal to these stably unmethylated Dam sites, we focus here on *ppiA* as in our data there are IPOD-HR occupancy changes at upstream Dam sites when *dam* is deleted (Figure 6CD). The *Δdam*-dependent loss in non-RNAP protein occupancy directly at Dam sites in a promoter region as seen with *ppiA* is precisely what we would expect to observe in a case of methylation-protein antagonism. However, it is not clear whether the *Δdam*-dependent occupancy change has an impact on transcription as there is a slight increase in RNAP-ChIP occupancy at the promoter in *Δdam* alongside a minimal – although statistically significant – decrease in *ppiA* transcript levels.

Integrating across all previous compatible data that we could identify (as detailed above), our RNA-seq data show the same directions of expression changes as prior studies for 13/15 cases (although the changes were not always statistically significant). We did not, however, observe evidence for local changes in protein occupancy at the promoters for most of those genes in response to methyltransferase deletion (with *ppiA* being the primary exception), indicating that either the identities of bound proteins change but the existence of binding does not, that the regulation due to the targeted methyltransferase is indirect, or that we are not sensitive in our assay to any changes that might occur.

### *dam* deletion mutants show loss of motility and downregulation of *flhDC*

One of the genomic loci with a dense clustering of 7 Dam sites was identified as the transcription start site of *flgN*, which encodes for a chaperone involved in cellular export of flagellum components^95,96^. Our RNAseq results show that *flgN* is downregulated in *dam* deletion mutants (Rockhopper mutants versus wild-type log_2_FC / q-values of *flgN*: −1.2 / 9.0 x 10^-16^ for *Δdam* vs. WT, −0.20 / 0.017 for *Δdcm* vs. WT, −0.97 / 2.9 x 10^-21^ for *Δdam/Δdcm* vs. WT). Additionally, there are mutant-specific IPOD-HR occupancy changes in the *flgN* promoter proximal to the Dam site cluster (Supplementary Figure 4). To our knowledge there is no experimental evidence for what regulatory proteins might act on the promoter immediately upstream of *flgN*, but two distal upstream promoters that impact *flgN* expression have previously been found to be occupied by CsgD and FlhDC^97–99^. While *csgD* transcript levels remain near-zero for all our genotypes, *flhC* and *flhD* transcript levels are decreased in our *Δdam* strains, and FlhDC was previously shown to activate expression of *flgN*^99^, thus providing a plausible path of information flow from *dam* deletion to decreased *flgN* transcription.

We next analyzed our datasets for information on the expression and protein occupancy of *flhDC*. While there is protein occupancy in the *flhDC* promoter, the occupancy pattern there appears to differ only marginally based on genotype, suggesting no major changes in the binding of regulatory factors upstream of *flhDC* (Figure 7AB). We examined the expression levels of the large set of known *flhDC* regulators across our genotypes to infer what regulators may be responsible for the occupancy signal in the *flhDC* promoter (Figure 7C). While this is an indirect inference, we also observed the mutant versus wild-type expression change across the regulons respective to each *flhDC* regulator to check which regulators of *flhDC* likely changed substantially in activity in each mutant (which would be indicated by the presence of changes in expression across the regulon of a factor that were coherent in sign with consideration of the effect of that regulator). The mutant versus wild-type log-fold change in expression of each regulon component was made positive if the change in expression matched the reported mode of regulation for the regulator-target pair or negative if the expression change opposed the annotated regulatory mode. The mean of these sign-changed log-fold expression changes within each regulon were then average to calculate the concerted log-fold change in expression of regulon for each regulator of *flhDC* (Figure 7D, Supplementary Figure 5), with a more positive change indicating stronger evidence for systematic changes throughout the regulon of a given upstream factor in line with its known regulatory effects.

**Figure 7:**
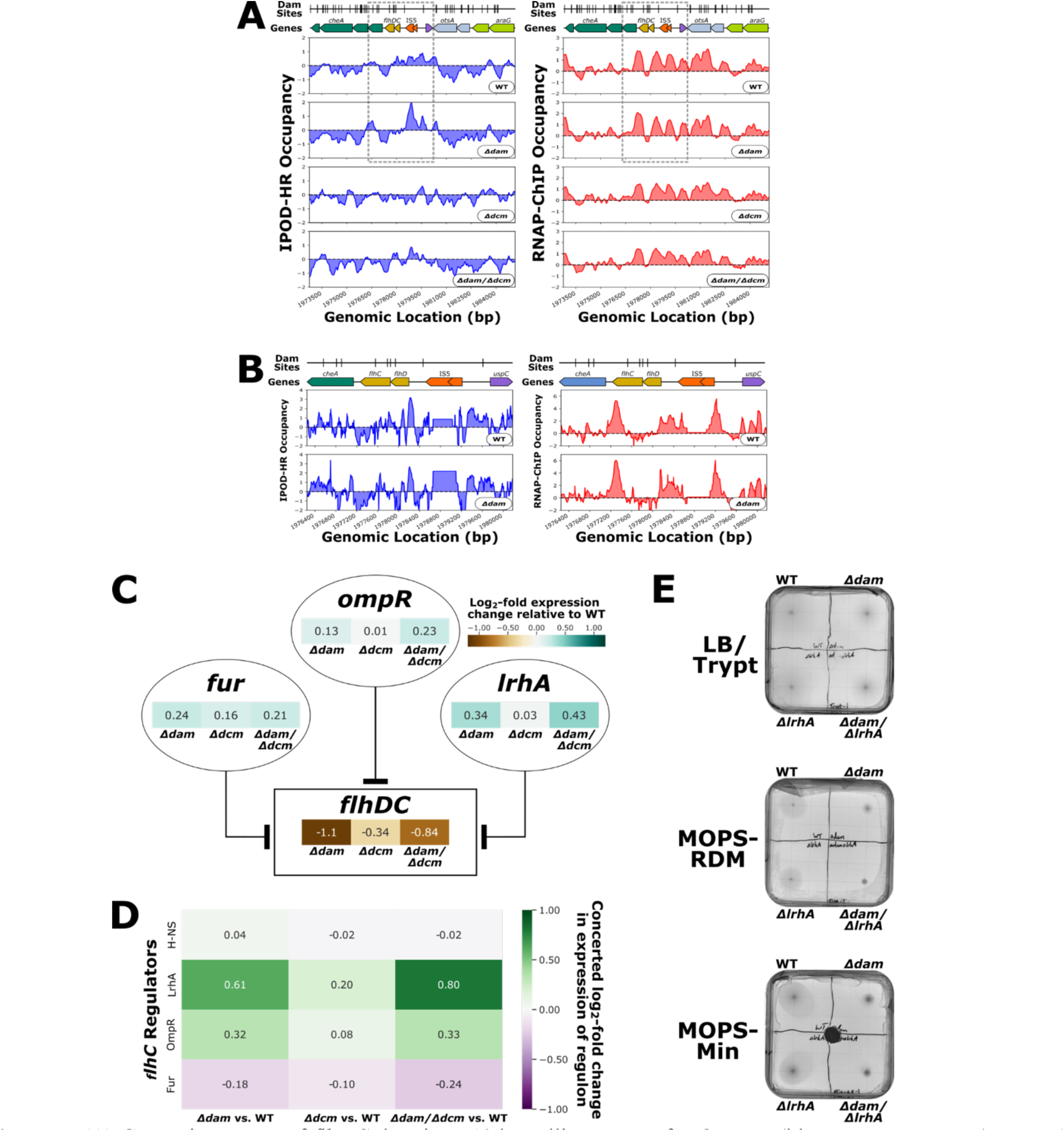
**(A)** Genomic context of *flhDC* showing 512 bp rolling mean of IPOD-HR (blue occupancy trace) or RNAP-ChIP (red occupancy trace) robust z-scores. Genes are differentially colored based on their membership to functional gene clusters. The dashed box designates the locus which is shown in panel B. **(B)** Immediate surroundings of *flhDC* (boxed region from panel **A**) showing IPOD-HR (blue occupancy trace) or RNAP-ChIP (red occupancy trace) robust z-scores. **(C)** Regulatory diagram including mutant versus wild-type log-fold expression change for selected subset of regulators of *flhDC*. The lines ending in flat bar arrowheads indicate repression. **(D)** Concerted log_2_-fold change in expression of the regulon of a selected subset of the regulators of *flhDC*. **(E)** Representative motility assay plates for LB/Tryptone, MOPS-RDM, and MOPS-Minimal conditions. The black circle in the center of the MOPS-Minimal plate is a filter disc soaked in aspartic acid which acts as a chemoattractant.

Of the known regulators of *flhDC,* LrhA stands out as having particularly high regulatory coherence scores in both of our strains lacking *dam* (Fig. 7D). LrhA is a repressor of *flhDC* transcription, and *lrhA* expression is increased in *Δdam* strains which is congruent with *flhDC* downregulation in *Δdam* genotypes^100^, and with the broader expression changes in the LrhA regulon in *dam* deletion mutants relative to WT. Fur and OmpR are also repressors of *flhDC*^101,102^ with increased expression in *Δdam* strains, but there is only a weaker degree of concerted log-fold change in OmpR regulon expression for *dam* deletion mutants relative to wild-type, and the Fur regulon does not show signs of strong concerted expression changes(Figure 7CD).

To characterize how loss of DNA methylation impacts flagellar motility, we performed swimming motility assays with wild-type, *Δdam, Δdcm*, and *Δdam/Δdcm* strains. To further characterize how loss of LrhA regulation of *flhDC* impacts motility relative to the motility impact associated with *dam* deletion, we additionally tested *ΔlrhA* and *Δdam/ΔlrhA* strains for swimming motility. While previous swimming motility assays have utilized tryptone-based motility plates^70^, the use of MOPS-RDM in our IPOD-HR and RNAseq procedures motivated us to develop MOPS-RDM and MOPS-glucose motility plates. Across all media types used for the motility assays, *dam* deletion mutants consistently demonstrated a substantial loss of motility as observed by the swimming distance of cells over time (Figure 7E), as would be expected based on our RNA-seq results. *lrhA* deletion mutants appear to have a small loss of motility in all tested media conditions, and the *Δdam/ΔlrhA* strain displays slightly less swimming motility than the *Δdam* strain. While we observe decreased LrhA expression and regulatory activity in *Δdam* strains, the swimming motility phenotypes (and particularly the lack of apparent epistasis between *dam* and *lrhA* deletions) do not provide further insight into the nature of the relationship between LrhA and Dam in regulation of motility; one thing that is clearly apparent is that the loss of motility of *Δdam* mutants cannot be attributed solely to the increase in LrhA activity, since the phenotype persists in the *Δdam/ΔlrhA* double knockout.

## DISCUSSION

Based on previous observations of a depletion of Dam methylation sites in extended, transcriptionally silent regions of high protein occupancy in the *E. coli* genome, we hypothesized that Dam methylation might play a global role in regulating the spread of NAP occupancy to control where EPODs occur, by inhibiting NAP occupancy in regions with relatively high Dam site densities. Contrary to our initial hypothesis, our results indicate that DNA methylation state (at least that arising from the native *E. coli* K12 DNA methyltransferases) does not substantially impact the global pattern of protein occupancy and EPOD formation at methylation sites. Thus, we observe that the genome-wide association between Dam sites and EPODs^46^ is not causal, but likely reflects other evolutionary constraints acting on long-term versus newly acquired genomic regions. Many EPODs in *E. coli* K-12 MG1655 have been observed to be associated with prophages and transposable elements^47^, which were likely acquired more recently and may have experienced less selective pressure, over less time, for containing Dam sites relative to more native regions of the genome^86,87,103^. While our data does not demonstrate a global pattern of DNA methylation state regulating local protein occupancy, there may still be a small number of methylation sites where antagonism exists between DNA methyltransferases and regulatory proteins. We also note that our tested conditions were limited to standard growth in rich medium (MOPS-RDM), and thus it is possible that changes in occupancy that would occur under other growth conditions are missed. For example, our cells were harvested at exponential phase but Dcm methylation appears to have a more biologically significant impact during stationary phase^22,36,104^. Another limitation of note is that deletion of methyltransferases is not expected to produce stably hemimethylated Dam sites which might more specifically interact with some DNA-binding proteins relative to unmethylated or fully methylated sites (as is the case with SeqA)^105–107^, and thus we might miss changes in occupancy arising specifically from the presence of hemimethylated regions.

Nevertheless, we did find that some protein occupancy changes caused by *dam* deletion are associated with loci featuring a dense clustering of methylation sites – such sites do show a decrease in total protein occupancy and an increase in RNA polymerase occupancy for *dam* deletion strains. Genomic regions with multiple proximal Dam sites have been previously identified as a potential regulatory element due to the poor processivity of Dam over such regions, resulting in hemimethylated sites^1,108,109^. Stably hemi- or unmethylated Dam sites are rare relative to fully methylated sites, and so they have been predicted to form specific associations with DNA-binding proteins such as SeqA^1,28^. However, loss of methylation at the dense Dam site clusters we identified does not appear to generally result in differential regulation of known local transcripts. We thus conclude that the *Δdam*-dependent changes in protein occupancy associated with dense clusters of Dam sites are primarily driven by the increased presence of RNA polymerase which is not transcriptionally active under our conditions, possibly due to increased promoter binding/transcriptional initiation without promoter clearance.

We observed the presence of an RNAP-ChIP peak proximal to all of our observations of high-density Dam site clusters at the ends of gene bodies, and we speculated that this RNAP-ChIP peak results either from direct recruitment of RNAP or stalling of RNAP at this site during Dam-associated mismatch repair^28,78,79^. Our samples for RNAP-ChIP and IPOD-HR are treated with rifampicin before crosslinking, and rifampicin inhibits promoter clearance of RNA polymerase^46,76,77^. Thus, we hypothesized that RNA polymerase recruited to an upstream promoter could read through a gene, get stalled at the Dam site cluster, and then be prevented from dissociating from the DNA by rifampicin until formaldehyde crosslinking. While our read end analysis did not support the presence of DNA damage at these loci, any potential direct signature of accumulations of strand breaks could easily have been masked by our sample workup, and we still suspect that stalling of the RNAP by DNA repair machinery is possible due to other aspects of dysregulated replication in *Δdam* strains such as asynchronous replication initiation and DNA base mismatches^80,110^. Contradicting the alternative scenario in which RNAP might be directly recruited to these Dam site clusters, we found a lack of changes in local transcript levels; however it is still possible that RNAP could be recruited for transcription of sRNAs^111^ but this transcription is not active (e.g. due to lack of promoter clearance) under our conditions, or that we were not able to detect these small transcripts.

To characterize another example of a dense Dam site cluster that shows substantial changes in protein occupancy, we considered a cluster occurring near the flagellar chaperone gene *flgN*. Our investigation was motivated by the presence of a dense Dam site cluster at the promoter of *flgN* as well as *Δdam*-associated downregulation of flagellum synthesis genes, which led us to characterize the regulatory network governing flagellar synthesis and swimming motility in our methyltransferase mutants. We focused on the master regulator FlhDC since it is a regulator of *flgN*, and *flhDC* expression is decreased in *Δdam* strains. To explore a possible causal relationship between DNA methylation and protein occupancy leading to *Δdam*-associated changes in the regulatory network governing flagellum synthesis, we expanded our investigation to include regulators of *flhDC*. Based on analysis of expression changes in the regulons of each *flhDC* regulator, we identified LrhA as the most likely regulator of *flhDC* to be differentially regulating its targets in response to methyltransferase deletion, but characterization of the swimming motility for *lrhA* and *dam* deletion strains did not reveal a clear regulatory relationship between LrhA and *dam* deletion. We also note that there are multiple transposable elements that may be incorporated upstream of *flhDC*, and the presence of these transposable elements has been shown to impact *flhDC* expression and flagellum-based motility^86,112^; we did not, however, observe any consistent pattern in our samples of changes in transposable elements around the *flhDC* promoter (data not shown). While the full nature of the relationship between DNA methylation and motility remains elusive, here we have demonstrated that loss of Dam methylation is associated with substantial loss of swimming-based motility.

While our observations suggest that loss of *lrhA* leads to a decrease in swimming motility, Lehnen *et al.*, 2002 transduced an insertionally inactivated *lrhA* into the MG1655 background and found that functional loss of *lrhA* leads to an increase in swimming motility^100^. Our laboratory strain of MG1655 has an IS1 insertion in the coding region of *dgcJ*^47,48^ which is a gene encoding for a diguanylate cyclase that has been associated with regulation of swimming motility^113^. It is thus possible that an epistatic interaction between *lrhA* and *dgcJ* explains the discrepancy in swimming motility phenotype resulting from functional loss of *lrhA* between our findings and those of Lehnen *et al*. 2002^100^, particularly given the importance of cyclic di-GMP for regulating flagellar motility^114^.

In globally surveying the impact of loss of DNA methylation on gene expression and protein occupancy in *E. coli* K-12 MG1655, our results indicate that although loss of *dam* and/or *dcm* leads to statistically and biologically significant changes in gene expression associated with observable phenotypes – such as loss of swimming motility – these changes appear to result primarily from global physiological effects of *dam* or *dcm* loss rather than being due to transcriptional regulatory consequences of losing local DNA methylation signal. Our observations of protein occupancy changes at methylation sites are primarily at loci with exceptionally dense clustering of Dam sites where we observe an increase in RNAP occupancy, but we find this pattern to be of no consequence to local transcriptional output. We thus conclude that DNA methylation is not a biologically significant factor in local gene expression or global chromatin structure for *E. coli* K-12 MG1655 under our tested conditions. Future studies that aim to address the question of whether there is any regulatory interplay between NAP or transcription factor occupancy and DNA methylation in MG1655 would be well-served by either testing a wider range of growth conditions or employing site-specific perturbation of methylation status without altering DNA sequence (e.g. with a tethered methyltransferase or demethylase)^115,116^.

## ACKNOWLEDGMENTS

This work was supported by NIH R35 GM128637 (to L.F.). H.A. was additionally supported by the University of Michigan Cellular and Molecular Biology Training Program (T32GM007315). The authors owe gratitude to Dr. Rebecca Hurto for technical assistance with the experiments described here, as well as Dr. Jeremy Schroeder for developing, and assisting in usage of, the IPOD-HR computational analysis pipeline. We also thank Amelia Lauth for running the IPOD-HR pipeline on our samples.

## SUPPLEMENTARY FIGURES

**Supplementary Figure 1:**
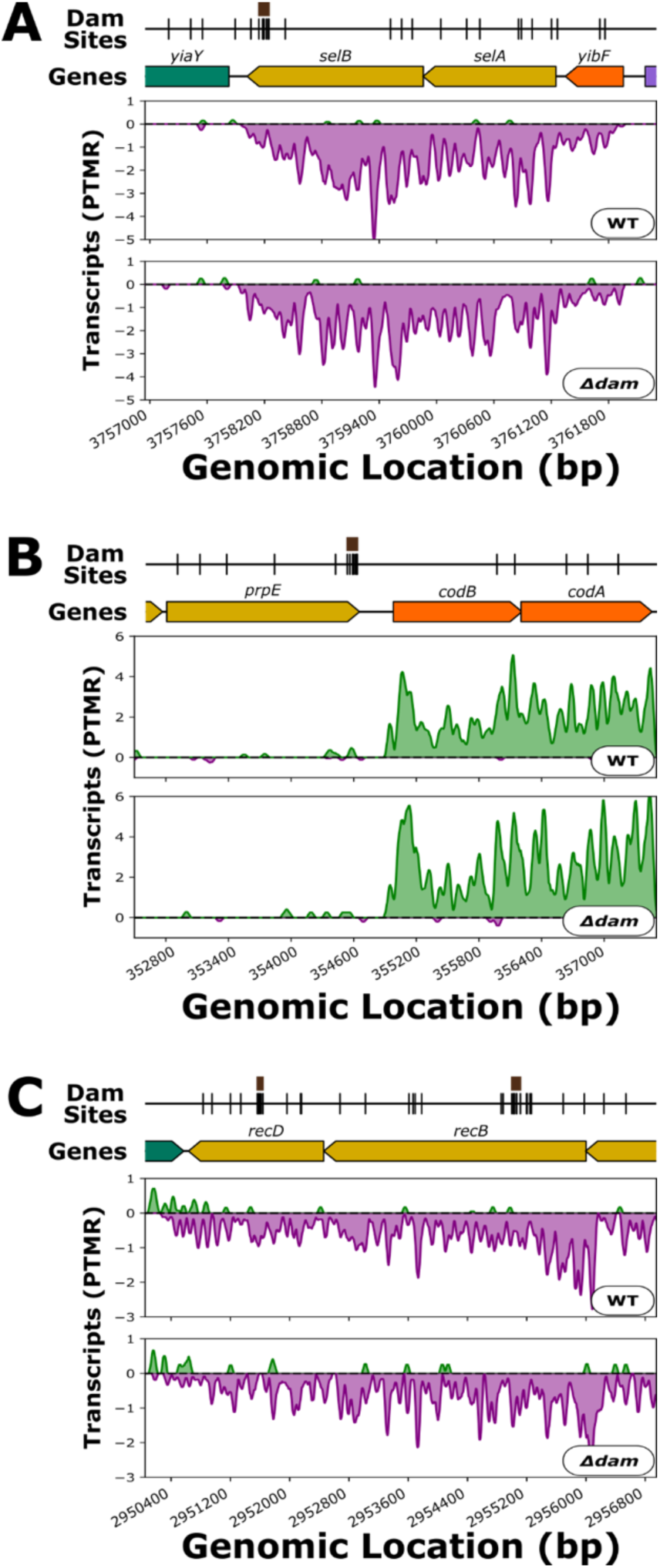
Genomic context of **(A)** *selB*, **(B)** *prpE*, and **(C)** *recBD* showing 51 bp rolling mean of RNAseq reads per tens of millions of reads (Transcripts (PTMR)) that were aligned to the positive (green occupancy trace) and negative (purple occupancy trace) strands. Brown boxes above markers on the “Dam Sites” tracks indicate “7 Dam Site Density” clusters of interest. Genes are differentially colored based on their membership to functional gene clusters.

**Supplementary Figure 2:**
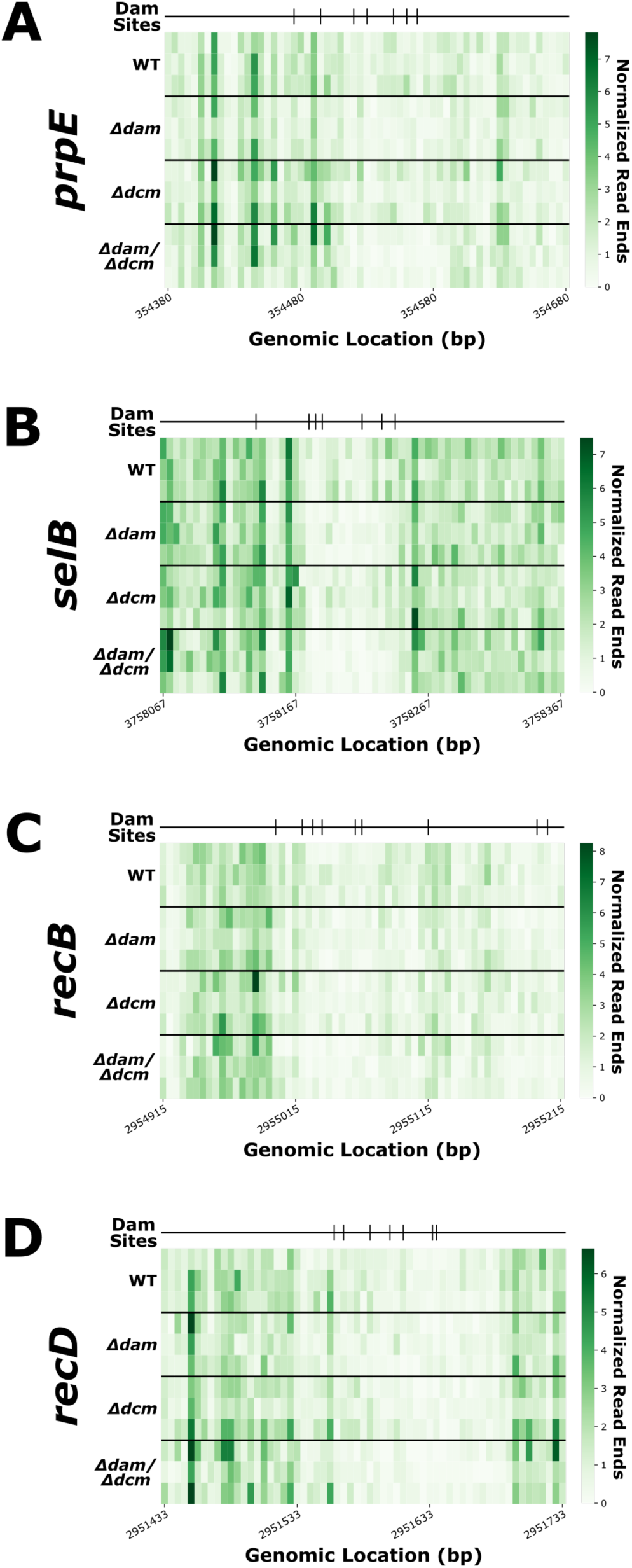
Normalized read ends, as calculated by the count of read ends at each genomic position divided by the total number of million read ends within each sample, around the “7 Dam Site Density” at the **(A)** *selB*, **(B)** *prpE*, **(C)** *recB*, and **(D)** *recD* loci in each of 3 replicates for each genotype.

**Supplementary Figure 3:**
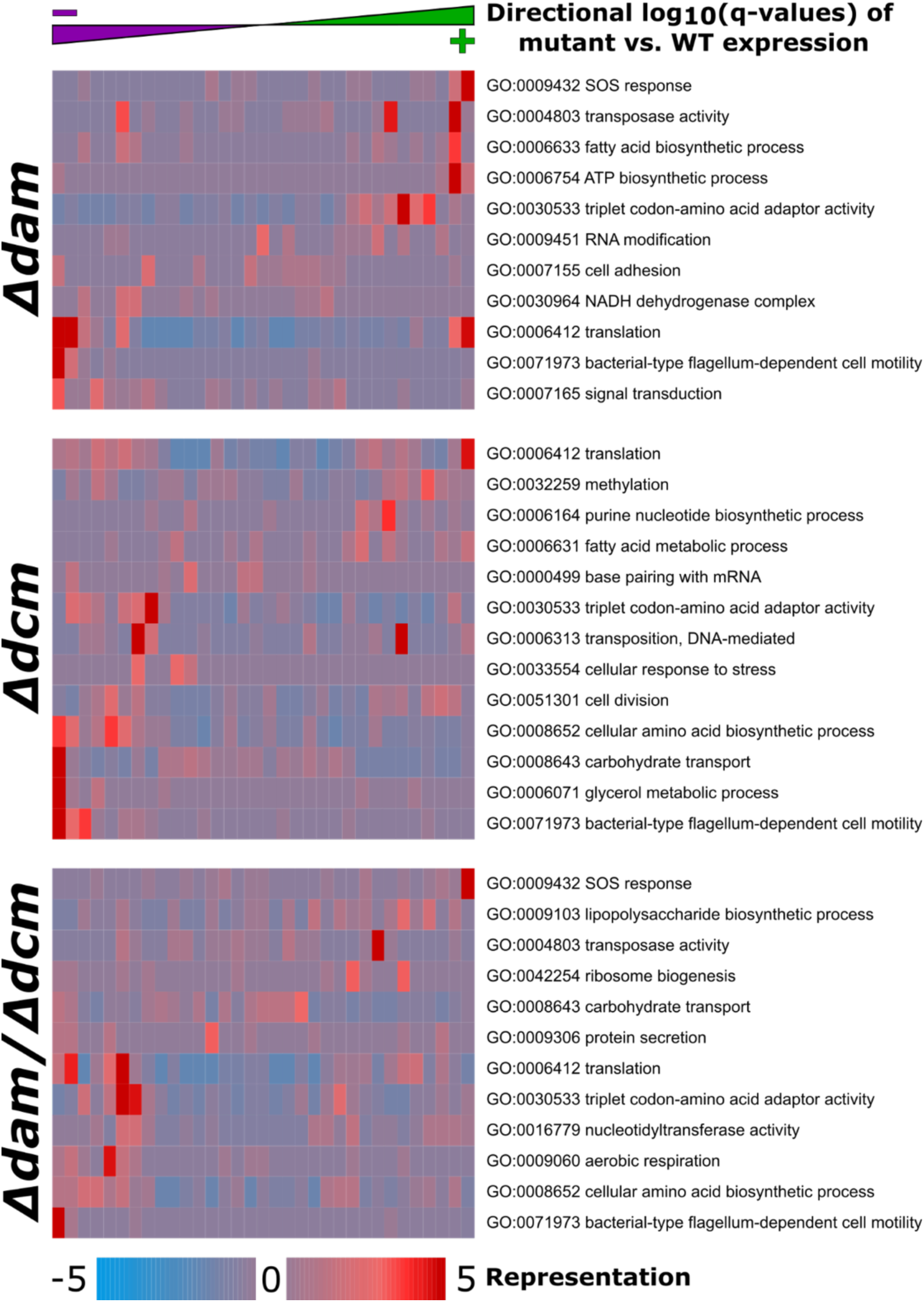
Gene set enrichment analysis for RNA-seq data across the indicated genotypes (relative to wild type). RNAseq data was analyzed using Rockhopper to produce q-values which assess statistical significance in expression change of each gene between strains. Directionality for expression change, where positive values indicate higher expression in the mutant relative to wild-type, was applied to the magnitudes of the log10(q-values). The values are divided into 21 evenly populated bins. iPAGE reports the representation of directional log10(q-values) across the genes annotated with each Gene Ontology (GO) term – thus, a redder bin indicates an over-representation of genes from the specified GO-term (row) at that expression change bracket (column).

**Supplementary Figure 4:**
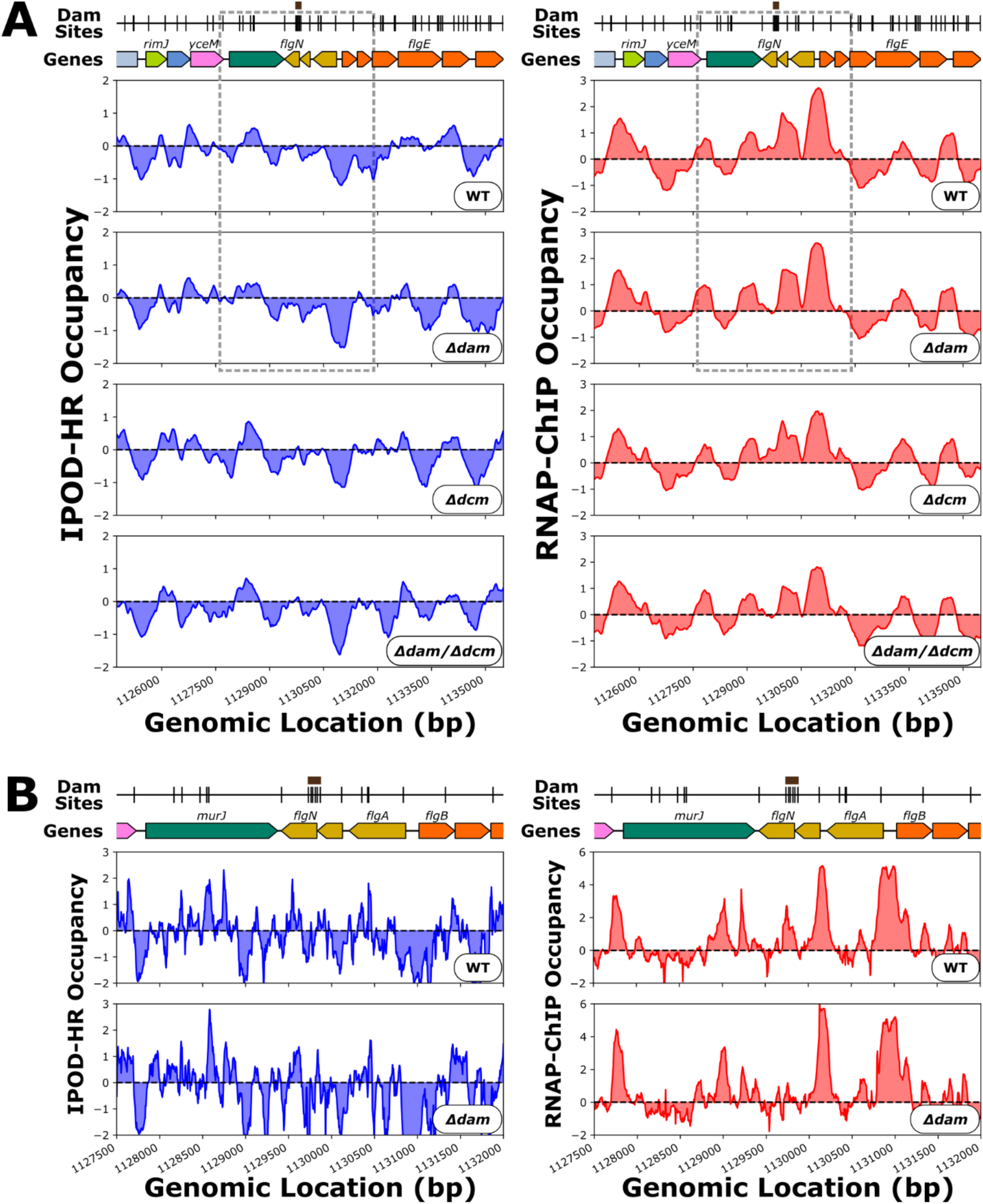
**(A)** Genomic context of *flgN* showing 512 bp rolling mean of IPOD-HR (blue occupancy trace) or RNAP-ChIP (red occupancy trace) robust z-scores. Brown boxes above markers on the “Dam Sites” tracks indicate “6 Dam Site Density” clusters of interest. Genes are differentially colored based on their membership to functional gene clusters. The dashed box designates the locus which is shown in panel B. **(B)** Genomic locus of *flgN* showing IPOD-HR (blue occupancy trace) or RNAP-ChIP (red occupancy trace) robust z-scores.

**Supplementary Figure 5:**
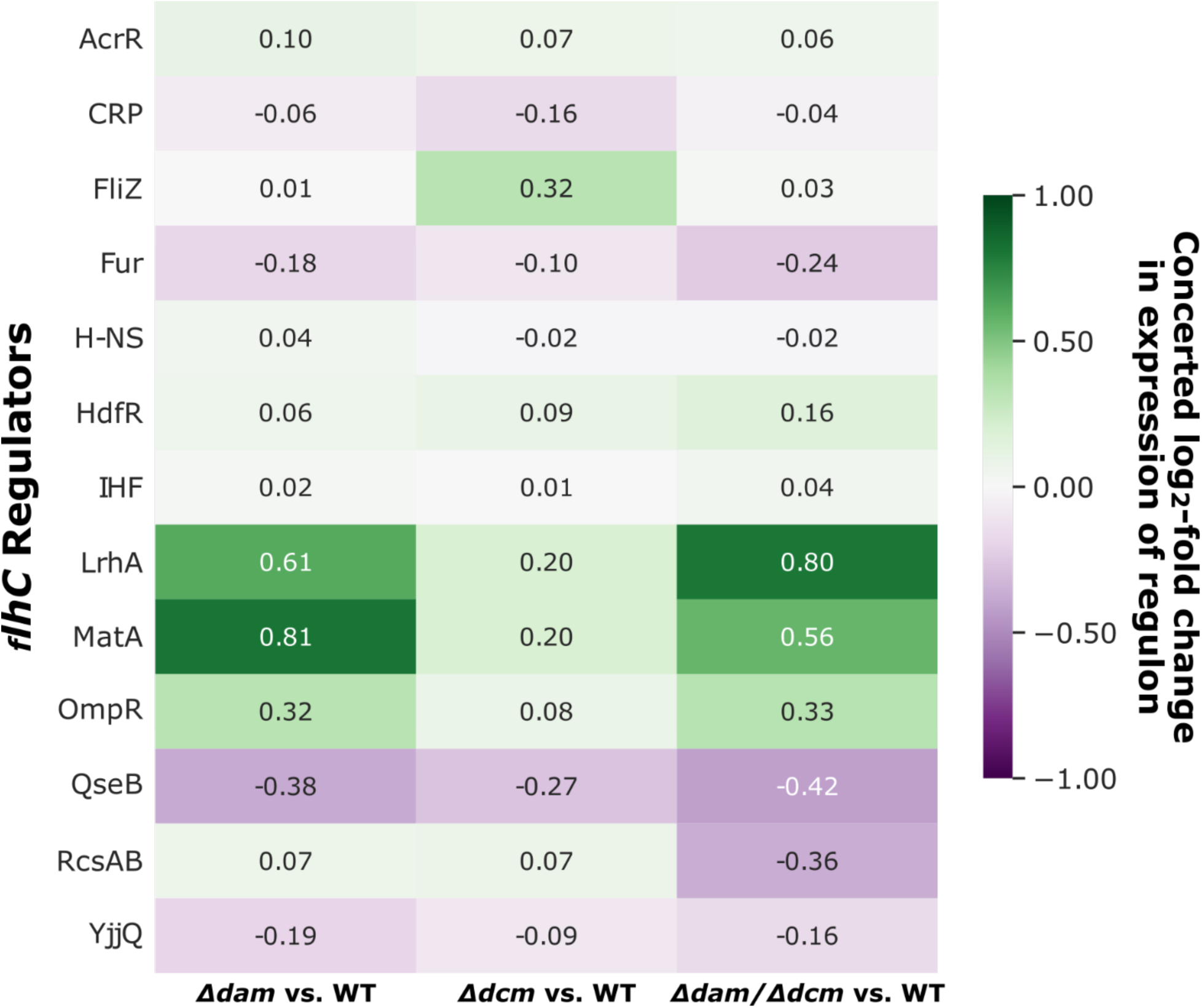
Heatmap representing the expression change in the regulon of regulators of *flhC*. We take the log ratio of mutant and wild-type expression values generated by Rockhopper for each gene in the regulon of the indicated *flhC* regulator. Directionality is then applied to determine whether expression changes in the regulon are consistent with the regulatory mode and expression of the regulator. E.g., genes that decrease in expression and are repressed by their regulator are “concerted” and thus contribute positively to the averaged log-fold change in expression.

